# Whole-genome comparisons identify repeated regulatory changes underlying convergent appendage evolution in diverse fish lineages

**DOI:** 10.1101/2023.01.30.526059

**Authors:** Heidi I. Chen, Yatish Turakhia, Gill Bejerano, David M. Kingsley

**Affiliations:** Department of Developmental Biology, Stanford University School of Medicine, CA; Department of Electrical and Computer Engineering, University of California, San Diego, San Diego, CA; Department of Biomedical Data Science, Stanford University School of Medicine, CA; Department of Computer Science, Stanford University School of Engineering, CA; Department of Pediatrics, Stanford University School of Medicine, CA; Howard Hughes Medical Institute, Stanford University, CA

## Abstract

Fins are major functional appendages of fish that have been repeatedly modified in different lineages. To search for genomic changes underlying natural fin diversity, we compared the genomes of 36 wild fish species that either have complete or reduced pelvic and caudal fins. We identify 1,614 genomic regions that are well-conserved in fin-complete species but missing from multiple fin-reduced lineages. Recurrent deletions of conserved sequences (CONDELs) in wild fin-reduced species are enriched for functions related to appendage development, suggesting that convergent fin reduction at the organismal level is associated with repeated genomic deletions near fin-appendage development genes. We used sequencing and functional enhancer assays to confirm that *PelA*, a *Pitx1* enhancer previously linked to recurrent pelvic loss in sticklebacks, has also been independently deleted and may have contributed to the fin morphology in distantly related pelvic-reduced species. We also identify a novel enhancer that is conserved in the majority of percomorphs, drives caudal fin expression in transgenic stickleback, is missing in tetraodontiform, *s*yngnathid, and synbranchid species with caudal fin reduction, and which alters caudal fin development when targeted by genome editing. Our study illustrates a general strategy for mapping phenotypes to genotypes across a tree of vertebrate species, and highlights notable new examples of regulatory genomic hotspots that have been used to evolve recurrent phenotypes during 100 million years of fish evolution.

## Introduction

Extensive efforts are underway to sequence the genomes of all eukaryotic species (Lewin *et al*., 2018, 2022) – including roughly 74,000 extant vertebrates (IUCN, 2022). Despite dramatic progress in the cost and throughput of whole-genome sequencing, it remains challenging to identify the genomic basis of interesting traits that have evolved repeatedly in wild species. The typical vertebrate genome spans several hundred megabases to several gigabases in length, of which only ~2% code for proteins (Biscotti *et al*., 2019; Gregory, 2022). Innovative new methods will be needed to compare and interpret genomes in order to identify both the protein-coding and the non-coding regulatory differences that underlie “endless forms most beautiful” that have evolved across the tree of life.

Fish species constitute nearly half of all vertebrates, and this enormous radiation exhibits especially remarkable phenotypic diversity (Norman, 1949; Nelson, Grande and Wilson, 2016; IUCN, 2022). A small handful of fishes have been developed as useful model organisms for studying vertebrate genetics and development, environmental monitoring, ecological interactions, and evolutionary change (Bell and Foster, 1994; Schier and Talbot, 2005; Ostlund-Nilsson, Mayer and Huntingford, 2006; Katsiadaki *et al*., 2007; Lleras-Forero, Winkler and Schulte-Merker, 2020; Patton, Zon and Langenau, 2021; Reid, Bell and Veeramah, 2021). Focused studies on a few models have been highly successful, but many more insights will likely come from comparative studies on thousands of additional fish species that have evolved in a diversity of environments.

A useful phenomenon within the teleost radiation includes the abundance of traits that have recurrently evolved across different clades. For example, changes in vertebral number, skeletal armor, teeth, scales, sensory modalities, locomotion, osmoregulation, temperature tolerance, and lifespan have evolved multiple times in independent lineages of fish (Norman, 1949; Nelson, Grande and Wilson, 2016; Kolora *et al*., 2021). Fin modifications – including extensive loss, dramatic expansion, and/or structural ornamentation – are particularly interesting given the outsized effects of fins on mobility, defense/predation, and reproductive success, and how easily fins can be scored visually or by non-destructive methods (Norman, 1949; Davenport, 1994; Westneat *et al*., 2004; Yamanoue, Setiamarga and Matsuura, 2010; Price, Friedman and Wainwright, 2015; Nelson, Grande and Wilson, 2016; Goldberg *et al*., 2019; Giammona, 2021; Sowersby *et al*., 2022). Because fins are homologous to tetrapod limbs, modification of these major body appendages in fish may also inform a variety of traits and diseases in other animals, including humans (Clack, 2009; Tanaka, 2016).

Recent studies suggest that recurrent evolution of phenotypic traits may often take place through reuse of particular genes (Conte *et al*., 2012; Martin and Orgogozo, 2013; Courtier-Orgogozo *et al*., 2020). As more species’ genomes are sequenced, it may therefore become possible to identify loci controlling certain traits by comparing phenotypes and genotypes over large phylogenetic trees where a trait of interest has evolved multiple times. This principle has previously been used to identify genomic loci involved in recurrent evolution of traits as diverse as vitamin C dependence, echolocation, antler loss, and flightlessness in birds (Hiller *et al*., 2012; Marcovitz *et al*., 2019; Sackton *et al*., 2019; Wang *et al*., 2019). Here, we use a growing number of sequenced fish species to identify genomic regions associated with recurrent fin modifications in fishes.

## Results

### A computational screen to identify genomic deletions recurrently associated with pelvic reduction

We selected 36 publicly available, full nuclear genome assemblies that pass a scaffold contiguity criterion of L50 ≤ 300 and that represent percomorph fish species informative for convergent pelvic fin evolution (Fig. 1A, see Supplementary File 1 for accession identifiers and sources). Most of the included species have complete bilateral pelvic fins and represent the ancestral, *outgroup* trait status (Fig. 1A-B) (Nelson, 1989). In addition, we included representative species from four independent *target* lineages that have evolved dramatic loss of pelvic appendages (Fig. 1A and 1C, Table S1). The two pelvic-loss clades that were best represented among the assemblies include three members of the family Syngnathidae (pipefish and seahorses) and four members of the order Tetraodontiformes (pufferfishes and Ocean Sunfish). Also included among our target species were one member of the family Synbranchidae (Rice Eel) and one member of the family Cynoglossidae (Tongue Sole, with one pelvic fin remaining on the blind side of the fish; see Table S1).

**Figure 1.**
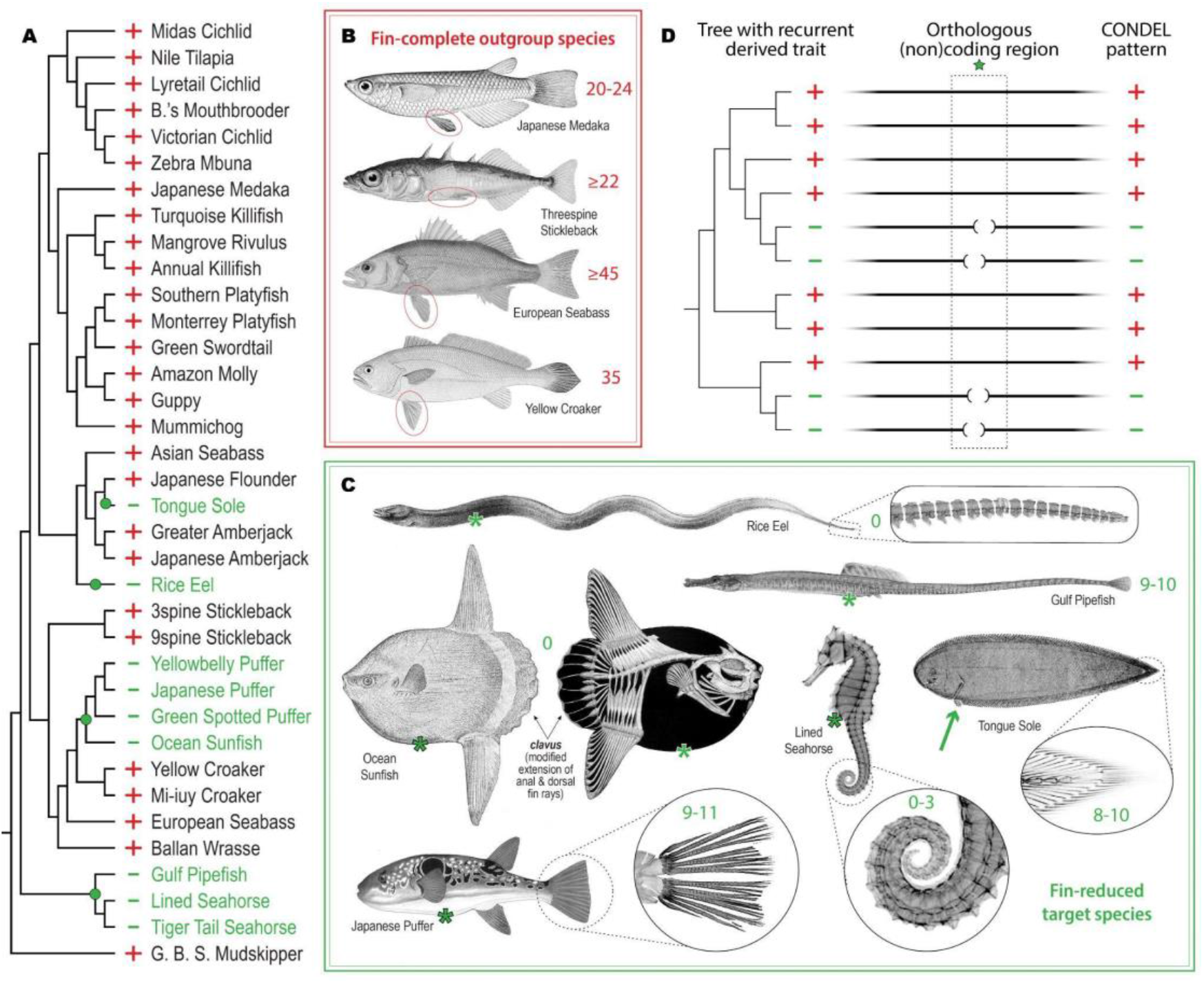
Genome-wide computational screen to identify conserved sequence deletions (CONDELs) associated with fin reduction. **A**, Pelvic and caudal fin reduction appears to have evolved at least four independent times (green dots) among the 36 percomorph species examined in this study. Fin-complete “outgroup” species and fin-reduced “target” lineages are denoted with a red plus (+) and green minus (-), respectively. **B**, Key outgroup species of the study include Japanese Medaka (the reference genome species and origin of the cell line used for *in-vitro* functional experiments), Threespine Stickleback (the species in which all *in-vivo* functional experiments were performed), and European Seabass and Yellow Croaker (the outgroups to which target lineages were compared in functional studies). All outgroup species exhibit the ancestral state of complete, bilateral pelvic fins (as indicated with red ovals) and of complete caudal fins supported by 20 or more bony rays (as indicated by the red numerical annotations adjacent to each tail fin). **C**, Representative species of target lineages exhibiting pelvic and caudal fin reduction include Tongue Sole (of Cynoglossidae); Rice Eel (of Synbranchidae); four members of the order Tetraodontiformes, including Japanese Puffer and Ocean Sunfish; and three members of the family Syngnathidae, including Gulf Pipefish and Lined Seahorse. With the exception of Tongue Sole (exhibiting one pelvic fin on the fish’s blind side, green arrow), all target lineages are pelvic-absent, as indicated by a green asterisk (*) positioned where pelvic appendages might *once* have existed in the ancestors of these lineages. Rice Eel, Ocean Sunfish, and the two seahorse species exhibit complete or near-complete caudal reduction, as indicated by three or fewer bony rays remaining at their caudal extremes. The tail fins of other target lineages exhibit no more than 11 caudal rays (indicated by the green numerical annotations adjacent to the caudal extreme of each species). See Table S1 and Table S6, respectively, for references documenting key species phenotypic status and for the sources of illustrations shown in **B** and **C**. **D**, The genome-wide association strategy implemented in this study requires a tree of broadly related species in which two or more independent sub-lineages have evolved a recurrent trait. Orthologous whole-genome alignments are then used to focus subsequent experimental interrogation on regions (green star) with tight correlation between genotype and phenotype – such as the regions of *<u>con</u>served sequence <u>del</u>etions* (CONDELs) associated with fin reduction identified here.

Based on overall genome assembly quality, extent of functional annotation, and outgroup trait status, we chose the Japanese Medaka genome assembly (abbreviated oryLat04) to be the reference against which all 35 other query genome sequences were aligned (Ichikawa *et al*., 2017). Using only orthology-confident alignment chains, we then searched for genomic sequences that were highly conserved in most outgroup species that had intact pelvic fins, but were missing (deleted or extensively diverged) in multiple syngnathid and tetraodontiform target species that showed pelvic loss (see Fig. 1D and Methods). We termed these regions *<u>p</u>ercomorph <u>con</u>served sequence <u>del</u>etions* or *pCONDELs* and reasoned that some subset of these candidate intervals are likely involved in the control of pelvic fin development. Because multiple species in Synbranchidae and Cynoglossidae were not able to be included, a genotype-phenotype match in these single-representative target clades was not strictly required in the computational screen, but was considered in selecting candidates for subsequent functional experiments.

With these criteria, we scanned intervals spanning 200 kb upstream to 200 kb downstream of the transcription start site for all successfully mapped reference gene orthologs. In total, we identified 1,614 predicted conserved sequences that were missing in multiple individuals of at least two independent target clades with pelvic reduction (see Methods). Of these pCONDELs, 9.5% intersected protein-coding exons, 49.6% intersected non-coding regions within genes, and 40.9% were located in intergenic regions of the Japanese Medaka reference assembly (see Supplementary File 2). Notably, the 3,489 gene orthologs that were linked to these candidate regions were most enriched for functions related to *medial fin development* (GO:0033338, 4.45-fold enrichment, *q*-value=0.022) and *embryonic appendage morphogenesis* (GO:0035113, 3.63-fold enrichment, *q*-value=0.026). These two most-enriched ontology terms suggest that convergent fin loss at the phenotypic level is associated with repeated genomic deletions that remove conserved sequences near fin and appendage development genes, including *Acvr1l*, *Bmp4*, *Dlx6a*, *Ext2*, *Extl3*, *Fgf10a*, *Fndc3a*, *Hmcn2*, *Lef1*, *Rspo2*, *Sall4*, *Shha*, *Smo*, *Sox9*, *Sp9*, *Tbx5a*, and *Wnt2ba* (see Supplementary Files 2 and 3 for the full list of genes linked to pCONDELs as well as all enriched ontology terms).

### Recurrent deletions at the *PelA* pelvic enhancer of *Pitx1*

One of the regions identified by the computational screen (pCONDEL.329) is a 50 bp interval that exhibits BLAST sequence similarity to the *PelA* pelvic enhancer of the limb and pituitary development gene *Pitx1*. In particular, pCONDEL.329 appears to be orthologous specifically to part of the minimal 500 bp core functional element of the *PelA* enhancer sequence (Fig. 2A) (Chan *et al*., 2010). Previous studies found that recurrent pelvic reduction in multiple freshwater populations of Threespine Stickleback is largely attributable to repeated, *de-novo* deletion of this non-coding pelvic control region (Shapiro *et al*., 2004; Chan *et al*., 2010; Xie *et al*., 2019). As in stickleback, the Japanese Medaka ortholog of the *PelA* enhancer is also upstream of the *Pitx1* gene. To confirm the presence of *PelA* deletions in other pelvic-reduced lineages, we amplified the orthologous region from relevant target species and their nearest outgroup (see Methods, Fig. 1–fig sup 1 and text sup 1, Supplementary File 4). Sanger sequencing confirmed that pelvic-reduced species show independent deletions with different sizes and breakpoints (see Methods, GenBank accession numbers in Table S2), resulting in loss of at least 78 bp of the 500 bp core *PelA* enhancer (corresponding to `gasAcu1-4.chrP:129,576-129,653` in the Threespine Stickleback reference genome).

**Figure 2.**
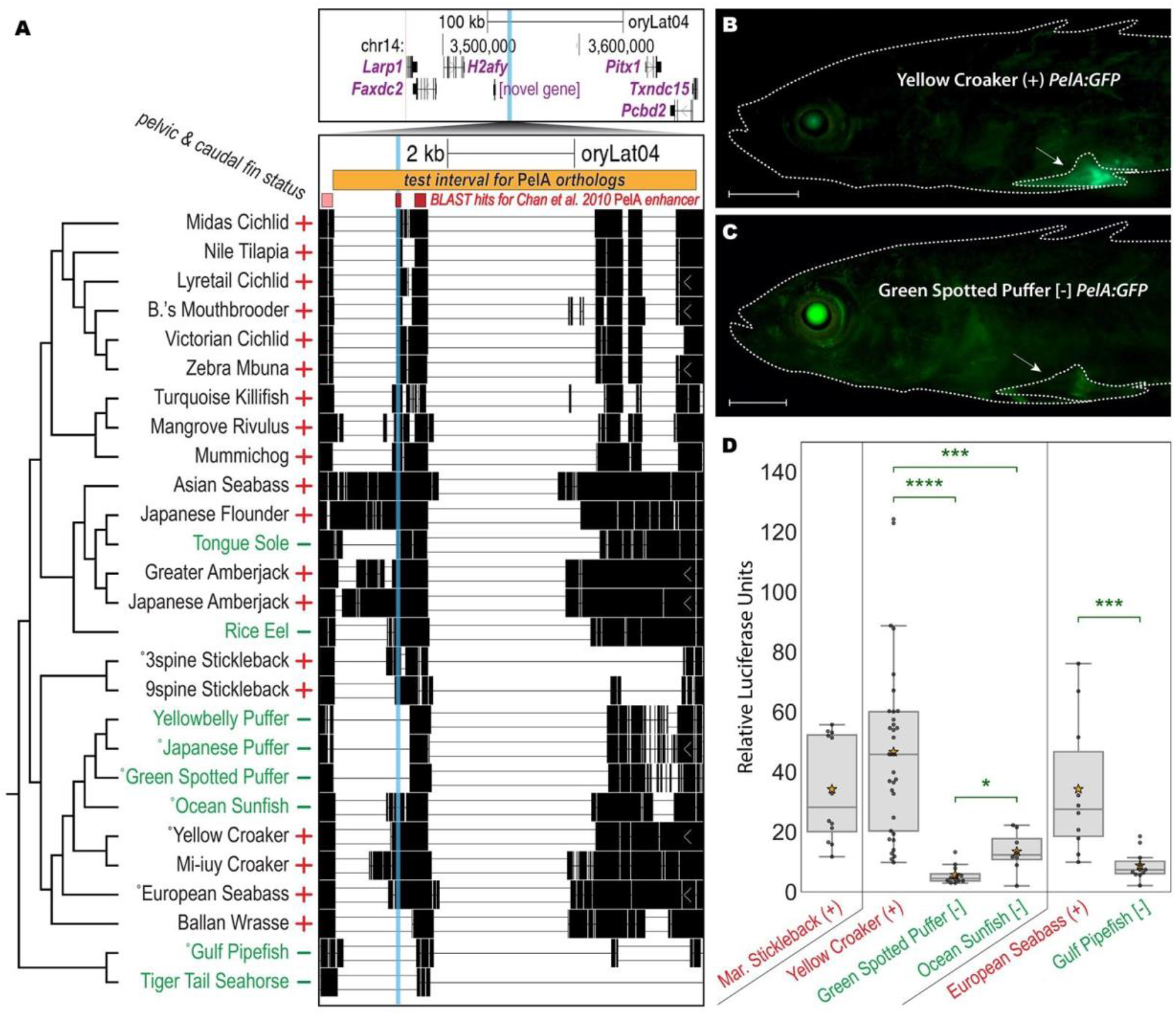
Recurrent deletions reduce functional activity of *PelA* enhancer sequences from multiple fish species. **A**, The 50 bp candidate interval pCONDEL.329 (cyan highlight) is located ~100 kb upstream of the hind appendage control gene *Pitx1* (top genome browser view), a region orthologous to *PelA* pelvic enhancer of the Threespine Stickleback *Pitx1* gene (Chan *et al*., 2010; Xie *et al*., 2019). BLAST similarity specifically to the 500 bp core *PelA* functional element is distinguished by the darker red bars and intersects the CONDEL interval. All existing pairwise orthologous alignment chains mapped to *Pitx1* are shown. The orange bar (`oryLat04.chr14:3,547,836-3,553,572`) marks the interval of orthologous sequence tested in the subsequent functional experiments depicted in **B-D** and/or verified by Sanger sequencing from the indicated (°) target and outgroup species. Red plus (+) and green minus (-) denote phenotypic trait status (presence or absence of pelvic and caudal fin reduction). **B-C**, GFP reporter expression in transgenic Threespine Stickleback driven by *PelA* orthologs from pelvic-complete (+) Yellow Croaker (**B**) and pelvic-absent [-] Green Spotted Puffer (**C**). Scale bars are 1 mm. White arrows point to the developing left pelvic girdle and spine. **D**, Relative luciferase expression in cultured fin cells driven by *PelA* orthologs from: pelvic-complete Marine Threespine Stickleback; pelvic-reduced Green Spotted Puffer and Ocean Sunfish and their nearest pelvic-complete outgroup Yellow Croaker; and pelvic-reduced Gulf Pipefish and its nearest pelvic-complete outgroup European Seabass. Boxes show quartiles; orange stars denote the mean of all points in the category. Two-tailed Mann Whitney U Test *p*-values: *, < 0.02; ***, < 4e-3; ****, < 1e-6.

To test whether these natural deletions also altered *PelA* function, we amplified the orthologous sequences denoted by the orange bar in Fig. 2A from several key species, cloned them upstream of a minimal promoter and GFP reporter gene, and injected the constructs into fertilized eggs from pelvic-complete Threespine Stickleback. The *PelA* sequence from outgroup Yellow Croaker drove bright GFP expression in developing pelvic structures of transgenic larvae (Fig. 2B, n=11/16 fish from 2 clutches), in patterns similar to that of the previously characterized marine stickleback ortholog (n=11/18 fish from 1 clutch, data not shown) (Chan *et al*., 2010). Only weak and diffuse pelvic expression was seen from the cloned *PelA* region of the pelvic-reduced Green Spotted Puffer (Fig. 2C, n=7/10 fish from 3 clutches), and this expression appeared substantially reduced compared to control eye expression known to occur from the *hsp70* minimal promoter of the transgenic construct (Fig. 2B-C) (Nagayoshi *et al*., 2008).

For a more quantitative assessment of the activity of various *PelA* orthologs, we cloned these regions upstream of a luciferase reporter gene and compared levels of luciferase activity following transfection into OLHNI-2 cultured fin cells derived from Japanese Medaka (Hirayama, Mitani and Watabe, 2006). Sequences from the previously characterized marine stickleback *PelA* region drove substantial expression in cultured fin cells, as did sequences from pelvic-complete outgroup species such as Yellow Croaker and European Sea Bass. In contrast, orthologs from two different pelvic-absent tetraodontiform species (Ocean Sunfish and Green Spotted Puffer), and from pelvic-absent Gulf Pipefish, drove significantly weaker luciferase activity compared to that of their nearest pelvic-complete outgroup (Fig. 2D). Previous studies have associated repeated deletions of *PelA* with recurrent pelvic reduction occurring over the last 10-20 thousand years of stickleback evolution (Shapiro *et al*., 2004; Chan *et al*., 2010; Xie *et al*., 2019). Our results identify similar deletions also operating over a significantly wider, 100+ million-year span of fish evolution, and confirm that our whole-genome comparative screen can identify functionally important lesions in known pelvic development loci.

### A novel, recurrently deleted regulatory region near *Faf1*

A previously unexplored genomic interval also highlighted by the computational screen is pCONDEL.1189, located in an intron of *Elavl4*. This 135 bp region appears to be either missing or disrupted in all scorable syngnathids and tetraodontiforms as well as in Rice Eel. We again confirmed the computational predictions in this region by amplifying and sequencing across the genomic interval pictured in Fig. 3A from relevant target species and their nearest outgroups (see Methods, GenBank accession numbers in Table S2).

**Figure 3.**
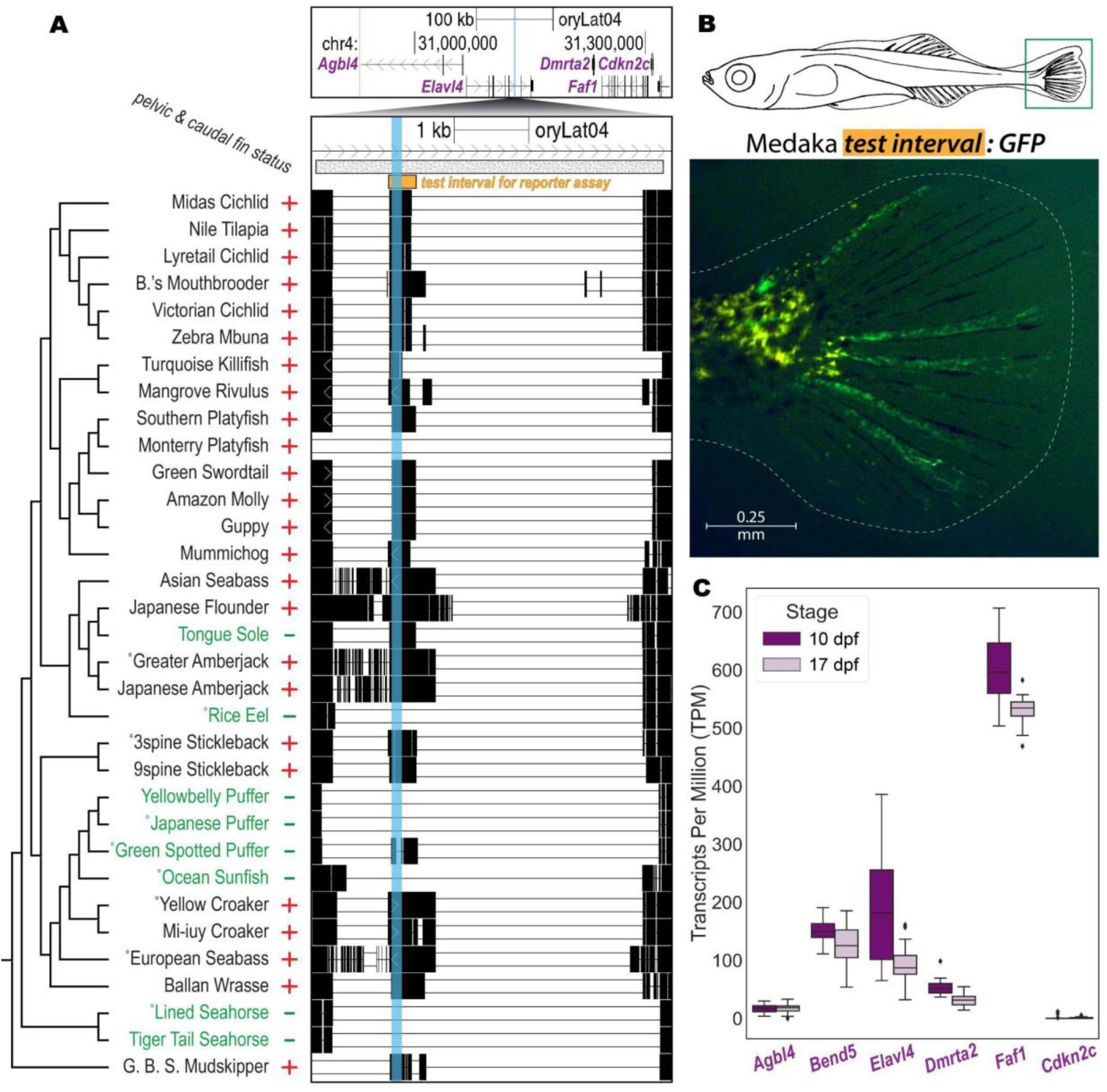
A novel, recurrently deleted regulatory region near *Faf1*. **A**, The 135 bp candidate pCONDEL.1189 (highlighted in cyan) is located within an intron of *Elavl4*. The genome browser view at top shows a region spanning 200 kb on both sides of pCONDEL.1189 (Japanese Medaka sequence space). Pairwise alignment chains for all species with orthologous mappings to *Elavl4* and *Faf1* are shown in the bottom genome browser view. The orange bar (`oryLat04.chr4:31,129,554-31,130,065`) marks the interval of orthologous sequence from Japanese Medaka tested in **B**. The hatched bar (`oryLat04.chr4:31,128,601-31,133,355`) marks the interval verified by Sanger sequencing from the indicated (°) target and outgroup species orthologs. Red plus (+) and green minus (-) denote phenotypic trait status (presence or absence of pelvic and caudal fin reduction). **B**, GFP reporter expression in the caudal fin (green box) of a transgenic Threespine Stickleback larva at Swarup stage 30 (schematic modified from Swarup 1958) driven by the orange test interval in **A** cloned from Japanese Medaka genomic DNA. **C**, Normalized gene expression data in the developing caudal fins of Threespine Stickleback (n=12 at each stage, before and after fin ray development) for all orthologous genes with transcription start sites predicted to lie within 200 kb of pCONDEL.1189 as shown in **A**. Data for *Bend5* – a gene between *Agbl4* and *Elavl4* found in Threespine Stickleback but not in Japanese Medaka – are also included. *dpf*, days post fertilization.

Given the absence of previous functional information about this candidate interval, we tested whether the conserved sequence element could drive informative GFP reporter expression in transgenic assays of enhancer activity *in vivo*. To do this, we cloned a ~500 bp test interval (orange bar in Fig. 3A) from Japanese Medaka genomic DNA upstream of a transposable GFP reporter and injected the construct into fertilized eggs of pelvic-complete Threespine Stickleback. Although we hypothesized that we would see enhancer activity in the pelvis of transgenic larvae, we instead observed consistent reporter expression in the developing caudal fin (Fig. 3B, n=15/21 transgenic fish from 6 clutches). These results suggested the candidate region might actually be a tail fin enhancer that was lost in many of the target lineages. A review of published literature and publicly accessible radiographs showed that the pelvic-reduced species in our study also exhibited varying degrees of reduction in this medial fin. Most pelvic-complete outgroup species showed totals of 20 or more segmented plus *un*segmented bony fin rays (lepidotrichia) present in their caudal fins (see numerical annotations in Fig. 1B and references in Table S1). In contrast, most pelvic-reduced species in our screen showed totals of 11 or fewer caudal fin rays (see numerical annotations in Fig. 1C). Note that all ecotypes of adult Threespine Stickleback possess 22 or more lepidotrichia (Fig. 1–fig sup 2 and see Plate III of the study published by Huxley in 1859), making this species an outgroup in terms of caudal fin status (Huxley, 1859; Lindsey, 1962).

Given the caudal fin enhancer activity of the pCONDEL.1189 region, we determined whether any genes in the surrounding genomic interval were also expressed in the developing caudal fins of Threespine Stickleback fry before and during lepidotrichia formation (10 and 17 days post fertilization, respectively) (Swarup, 1958). Of the genes with transcription start sites within 200 kb of pCONDEL.1189 in Japanese Medaka genomic space, *FAS-associated factor 1* (*Faf1*) was most highly expressed (Fig. 3C). *Faf1* encodes a protein that associates with the cell death receptor FAS (TNFRSF6), and has been shown to modulate apoptosis and cell proliferation, including in skeletal tissues (see Discussion).

### Morphological effects of engineered mutations in pCONDEL.1189 and *Faf1*

To test whether pCONDEL.1189 was required for normal caudal fin development, we used CRISPR-Cas9 editing in Threespine Stickleback to remove the conserved sequence from an outgroup species that normally exhibits complete caudal and pelvic fin development (Fig. 4A). Following injections of Cas9 and sgRNAs into fertilized stickleback eggs, genotyping and sequencing confirmed high efficiency removal of the conserved element in resulting fish (Fig. 4–fig sup 1A-C). Across 17 genome-edited clutches, 8% of the pCONDEL.1189-targeted fish (n=14/175) exhibited ectopic overgrowth of the typically unsegmented procurrent rays on the dorsal edge of the tail (Fig. 4B-C; Fig. 4–fig sup 1D; and Fig. 4–fig sup 2). To control for potentially confounding effects of Cas9 nuclease injection at the zygote stage, we simultaneously injected clutch siblings with equal concentrations of editing reagents (see Table S3) that instead targeted the previously characterized *Slc24a5* gene of the *golden* pigment-control locus (Lamason *et al*., 2005; Wucherpfennig, Miller and Kingsley, 2019). Only a single *golden* knockout control sibling also showed tail abnormalities (0.43%, n=1/236, Fig. 4–fig sup 2), confirming a significant effect of pCONDEL.1189 targeting on caudal fin development (*p*=3.55e-5).

**Figure 4.**
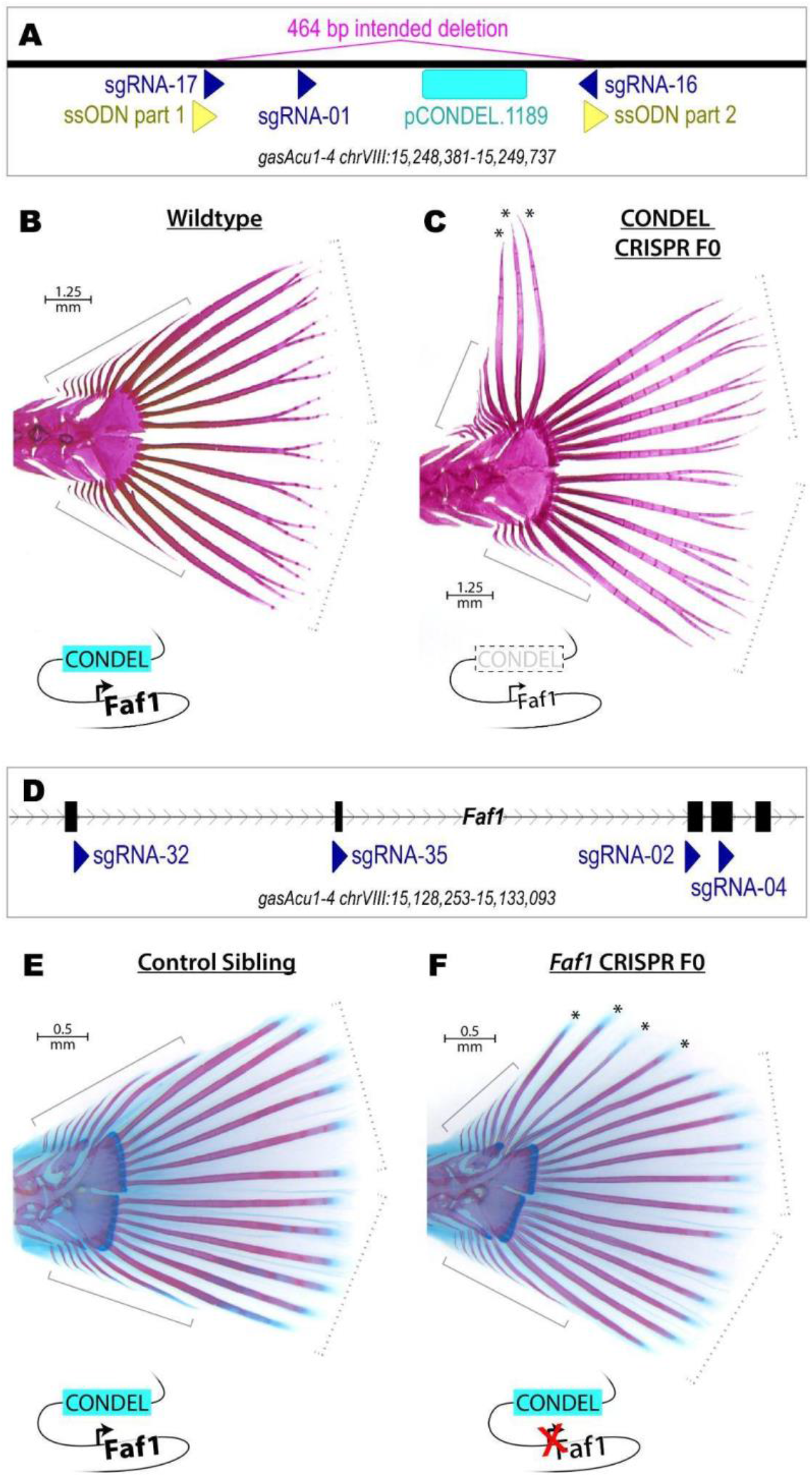
Morphological effects of engineered mutations in pCONDEL.1189 and *Faf1*. **A**, The conserved element at pCONDEL.1189 (cyan bar) was deleted from Threespine Stickleback by injecting zygotes with Cas9 protein, 3 sgRNAs (dark blue triangles), and a 60 bp single-stranded DNA oligonucleotide (a concatemer of the two yellow triangles) to promote homology directed repair and creation of the indicated 464 bp deletion. **B**, Alizarin red-stained wildtype Threespine Stickleback exhibiting the typical 6 segmented *principal* rays each on the dorsal and ventral halves of the tail fin (dashed brackets) as well as a variable number of smaller, unsegmented *simple* rays more anteriorly (solid brackets). **C**, Alizarin red-stained mosaic F0 crispant targeted for deletion at the pCONDEL.1189 region showing additional long segmented rays (*) where typically only unsegmented simple rays exist. **D**, *Faf1*, putative target gene of the conserved element enhancer at pCONDEL.1189, was disrupted by injecting Threespine Stickleback zygotes with Cas9 protein and 4 sgRNAs (dark blue triangles) targeting exons 2-5 of the gene (furthest right exon 6 is also pictured). **E**, Alizarin red and alcian blue-stained unmodified control sibling exhibiting the typical 6 segmented *principal* rays each on the dorsal and ventral halves of the tail fin (dashed brackets) as well as a variable number of smaller, unsegmented *simple* rays more anteriorly (solid brackets). **F**, Alizarin red and alcian blue-stained fish targeted for inactivation of *Faf1* showing ectopic segmented rays (*) where typically only unsegmented simple rays exist. Schematics summarizing the genome modification performed near pCONDEL.1189 and *Faf1* are included below **B-C** and **E-F**.

To test whether the hypothesized target gene, *Faf1*, was similarly required for normal caudal fin development, we also injected Threespine Stickleback zygotes with a pool of 4 sgRNAs targeting early exons of the gene (Fig. 4D). Sequencing confirmed the creation of indels disrupting *Faf1* coding exons in resulting fish (Fig. 4–fig sup 3A-C). Across 10 genome-edited clutches, we observed that 6% of fish (n=16/265) exhibited a phenotype very similar to that of the enhancer deletion fish, with unusually long and segmented rays on the dorsal edge of the tail (Fig. 4E-F; Fig. 4–fig sup 3D; and Fig. 4–fig sup 4). No mutant tail fin phenotypes were seen in 277 unmodified control siblings from the same clutches, confirming a significant effect of *Faf1* targeting on caudal fin development (*p*=6.66e-6).

## Discussion

Previous studies have identified “hotspots of evolution” — particular genomic regions that are used repeatedly when similar traits evolved in multiple lineages (Martin and Orgogozo, 2013). Over 115 hotspot loci are known to have been repeatedly used during independent evolution of traits across closely related populations within a species (intraspecific hotspots). In contrast, about four-fold fewer loci are currently known to underlie repeated evolution at higher taxonomic levels (i.e. hotspots reused between genera and even more distant phylogenetic taxa) (Courtier-Orgogozo *et al*., 2020).

Extensive linkage mapping, mutation, transgenic, and genome-editing studies have previously shown that the *PelA* enhancer of the *Pitx1* gene corresponds to an intraspecific genomic hotspot for repeated pelvic loss within Threespine Stickleback (*Gasterosteus aculeatus*) (Shapiro *et al*., 2004; Chan *et al*., 2010; Xie *et al*., 2019). Multiple *Gasterosteus* populations have independently evolved total loss of pelvic fins in new freshwater lakes generated by widespread melting of glaciers in the last 20,000 years. The mutation spectrum in these pelvic-reduced *Gasterosteus* populations consists of deletions of a few hundred to a few thousand bases that completely eliminate a shared 369 bp region of the *PelA* pelvic enhancer control region (Chan *et al*., 2010; Xie *et al*., 2019). Pelvic reduction can be phenocopied by targeted deletions of the *PelA* region (Xie *et al*., 2019), or rescued by reintroducing *PelA:Pitx1* constructs into pelvic-less fish (Chan *et al*., 2010), confirming that *PelA* is a key causal locus for loss of pelvic structures in *Gasterosteus*.

Genetic crosses also suggest that changes in the *Pitx1* locus underlie some examples of pelvic reduction in the more distantly related Ninespine Stickleback lineage (*Pungitius pungitius*) (Shapiro *et al*., 2009; Shikano *et al*., 2013; Kemppainen *et al*., 2021). However, the genetic basis of pelvic reduction appears to be heterogeneous across different *Pungitius* populations, and the causative molecular lesions in most populations are still unknown (Shapiro *et al*., 2009; Shikano *et al*., 2013; Kemppainen *et al*., 2021).

Our current study has identified recurrent deletions of the *PelA* enhancer occurring in distantly related wild fish species that have independently evolved pelvic reduction since their divergence over 100 million years ago. These results show that the *PelA* locus is not only an intraspecific evolutionary hotspot, but also an interordinal hotspot that is reused when pelvic reduction evolves in many different lineages. Most previous examples of hotspot loci reused between distant phylogenetic groups have involved repeated amino acid changes in particular proteins (Courtier-Orgogozo *et al*., 2020). *PelA* is one of a single-digit number of *cis*-regulatory regions now known to underlie naturally evolved convergent traits spanning intergeneric distances and above (Sagai *et al*., 2004; Miller *et al*., 2007; Guerreiro *et al*., 2013; Kvon *et al*., 2016; Wucherpfennig *et al*., 2022).

Although the *PelA* region had previously been implicated in pelvic reduction, most of the other genomic regions recovered in our genome-wide comparative analysis do not have previously known functions. Over 90% map outside of protein-coding exons of genes, so, like *PelA*, may also correspond to regulatory elements in the genome. The genomic regions near these recurrently lost sequences are enriched for genes involved in fin and appendage development, and we hypothesize that regulatory deletions occurring near such genes are an important contributor to recurrent fin evolution in fishes.

Our studies clearly illustrate the value of additional experimental follow-up on novel loci identified by comparative genomic scans. Although we selected target and outgroup species to screen for regions associated with pelvic reduction, when we tested the *in-vivo* activity of the conserved element at pCONDEL.1189, we found that this non-coding regulatory region drove expression in caudal fins rather than pelvic fins of transgenic larvae. Retrospective trait inspection showed that pelvic *and* caudal fin reduction are correlated with each other in the original species tree. Thus, the genomic regions recovered in our whole-genome screen might be associated with either of these fin phenotypes (or with any other trait that co-occurs with pelvic and caudal fin reduction in the same target lineages).

Our genome editing experiments further confirm that the pCONDEL.1189 region functions during caudal fin development. Deletion of the region results in the formation of additional long segmented fin rays in the caudal fins of Threespine Stickleback. Targeting of the nearby *Faf1* gene produces a very similar phenotype, suggesting the pCONDEL.1189 region likely acts by regulating *Faf1* function. *Faf1* is named for its association with the *Fas1* receptor, which activates cell death pathways in response to intercellular signaling (Chu, Niu and Williams, 1995; Ryu *et al*., 2003). The extra fin rays seen in our targeting experiments may thus be the result of decreasing the activity of a known apoptotic pathway during caudal fin development.

It is interesting that tail phenotypes are only observed in a fraction of CRISPR/Cas9-edited founder fish (approximately 8% of pCONDEL.1189-targeted fish, and 6% of *Faf1*-targeted, Fig. 4—fig sup 1D and 3D). In addition, we have not observed similar tail phenotypes when we intercrossed founders to produce F1 fish that are homozygous for mutant pCONDEL.1189 alleles in every cell during normal development (Fig. 4—fig sup 5). These results might be attributable to the polar coordinate model of limb development first described by French and colleagues (French, Bryant and Bryant, 1976). Based on observations of supernumerary limb and digit formation in insects and amphibians after mirror-image appendage grafting, this model posits that every cell has a particular positional identity relative to all other cells in a given tissue, as if arranged in a polar coordinate system. When normally non-adjacent identities come into contact, either by injury, tissue grafting, or – we suggest – mosaic genome editing, cells proliferate and/or intercalate to restore the original cell-cell adjacencies and thereby generate supernumerary structures. We hypothesize that the ectopic fin ray phenotype in our mosaic founders may occur when *Faf1* activity is disrupted in subsets of developing fin cells, and perhaps only when those cells occur in juxtaposition with other cells that have normal *Faf1* activity. Future studies could modulate *Fas1/Faf1* activity in subgroups of cells during caudal fin development to test whether manipulation of this pathway in specific anatomical regions leads to predictable morphology that resembles fin ray changes in wild fishes.

Our current work with *PelA* and pCONDEL.1189 illustrates how genome-wide approaches can identify both known and novel loci contributing to repeated evolution of fish fin modifications. In our genomic comparisons, we focused on a specific type of convergent DNA change (deletions), because such structural changes seemed likely to yield phenotypic consequences, and were already associated with repeated evolution of pelvic reduction in multiple stickleback populations (Chan *et al*., 2010; Xie *et al*., 2019). However, we note that similar genome-wide association approaches can be designed to identify many other types of DNA alterations, including convergent amino acid changes, point mutations, splicing changes, insertions, copy number differences, or multiple types of lesions co-occurring in the same gene or gene pathway in different lineages (Hiller *et al*., 2012; Hiller, Schaar and Bejerano, 2012; Chikina, Robinson and Clark, 2016; Marcovitz, Jia and Bejerano, 2016; Prudent *et al*., 2016; Lowe *et al*., 2017; Partha *et al*., 2017; Berger *et al*., 2018; Marcovitz *et al*., 2019; Sackton *et al*., 2019; He *et al*., 2020; Kowalczyk *et al*., 2020; Turakhia *et al*., 2020; Roscito *et al*., 2022; Kaplow *et al*., 2022; Kowalczyk, Chikina and Clark, 2022).

It is still not clear how often repeated phenotypic evolution is due to convergent mutational changes in the same genomic regions (Gompel and Prud’homme, 2009; Conte *et al*., 2012; Martin and Orgogozo, 2013). Recent reviews of known cases suggest the answer may vary depending on the evolutionary distance among lineages in which convergent phenotypes arise, with more closely related species being most likely to evolve through similar genetic pathways (Conte *et al*., 2012; Ord and Summers, 2015). On the other hand, reuse of certain genomic loci still appears to be appreciable even among distantly related lineages (Conte *et al*., 2012; Martin and Orgogozo, 2013; Courtier-Orgogozo *et al*., 2020), and our results provide striking new examples of genomic convergence occurring across distantly related fish species that last shared a common ancestor over 100 million years ago. Although we focused on fin modifications in this study, many other types of traits also evolve repeatedly when different lineages adapt to similar ecological pressures (McGhee, 2011). As fish make up nearly half of vertebrate species and also account for a large fraction of known examples of repeated evolution (Ord and Summers, 2015), they may provide an especially powerful system for linking phenotypes and genotypes using tree-wide association and functional genomic studies.

## Materials and Methods

### Computational screen to identify conserved sequence deletions (CONDELs)

#### Genome assemblies

See Supplementary File 1 for accession identifiers and sources.

#### Repeat masking

For assemblies `hipEre01` and `synSco01, RepeatMasker v4.1.0` (with NCBI/RMBLAST [`2.10.0`+] and `Complete Master RepeatMasker DatabaseCONS-Dfam_3.1-rb20181026`) were used to soft-mask repetitive sequences prior to whole-genome alignment (Smit, Hubley and Green, 2015). Specifically, the following commands were run:

~~~
RepeatMasker -engine rmblast -species ‘hippocampus erectus’ -s -no_is -cutoff 255 - frag 20000 hipEre01.fa
RepeatMasker -engine rmblast -species ‘syngnathus scovelli’ -s -no_is -cutoff 255 - frag 20000 synSco01.fa
~~~

#### Whole-genome alignment and orthology mapping

Based on its high level of contiguity and functional annotation, the Japanese Medaka genome assembly ASM223467v1 (Genbank accession # GCA_002234675.1, assembly abbreviation `oryLat04`) was used as the reference to which all 35 other query genome sequences were aligned (Ichikawa *et al*., 2017). Ensembl release 98 gene models for ASM223467v1 served as the genomic landmarks for identifying orthologous alignment chains (Cunningham *et al*., 2022). BLASTZ-based pairwise whole-genome alignment chains (Kent *et al*., 2003) were generated using the doBlastzChainNet.pl tool (https://github.com/ENCODE-DCC/kentUtils/, last accessed 12 Sep 2022) – with the `chainLinearGap` setting as `medium` – and the parameters listed in Supplementary File 1. Only alignment chains containing confident gene ortholog mappings, identified as previously described (Turakhia *et al*., 2020) with `gene-in-synteny` and `second-best-chain-ratio` thresholds of 10, were used in subsequent computational steps. All orthologous pairwise alignment chains can be viewed at the following UCSC Genome Browser assembly hub (Kent *et al*., 2002; Raney *et al*., 2014): https://genome.ucsc.edu/s/hchen17/oryLat04_public.

#### Identifying conserved sequences and gaps in alignment chains

Orthologous pairwise alignment chains allow for convenient programmatic distinction between reference genome regions that are either intact and conserved at the orthologous position of the aligned query genome versus those that are missing or extensively diverged (Kent *et al*., 2003). We thus used the chains with gene ortholog mappings from the previous step to record single- and double-sided chain gaps (intervals predicted to be missing from the aligned genome relative to the reference) from each target species. To avoid mistaking incomplete genome assembly for genuinely missing sequence, we excluded (subtracted) chain gap intervals that are within 100 bp of an assembly gap longer than [N]_5_ in any given target genome. For each alignment chain, we merged any chain gap intervals that are within 20 bp of each other.

Using orthologous chains of outgroup species, we identified *conserved* genomic intervals by (a) recording regions in each query genome that exhibit an intact alignment block; (b) computing percent sequence identity relative to the `oryLat04` reference using 10 bp, 25 bp, 50 bp, and 100 bp sliding windows across the alignment blocks; (c) sorting the windows by percent sequence identity; and (d) keeping the top *M* most conserved windows that, when merged and flattened, cover 5% of `oryLat04` and together were deemed per-species *conserved elements*. For each alignment chain, we merged conserved elements within 20 bp of each other. Note that the conserved elements encompass all additional windows with sequence identity scores tied with that of the *M*^th^ (lowest-scoring) window included to reach the 5% reference coverage threshold. Summary statistics for the sliding windows used to delineate conserved elements are in Table S4.

#### Screen for genomic regions associated with fin reduction

For each Japanese Medaka reference gene that was successfully mapped to at least 17 pelvis-complete outgroups and 5 pelvis-reduced target species, we scanned the interval spanning 200 kb upstream and downstream of the canonical (longest) isoform’s transcription start site (400,001 bp total). Within this interval, we identified *target deletions* by recording regions where sequence is missing (i.e. a valid chain gap is present) in at least ⅔ of all species in each of the two officially screened target clades (Tetraodontiformes and Syngnathidae). We then intersected the target deletions with regions covered by conserved elements in at least 17 outgroups (not including the reference assembly itself, which is sequence-conserved by definition) and spanned at least 20 bp to identify raw candidate conserved sequence deletions (CONDELs). We merged raw CONDEL candidates within 20 bp of each other and only recorded final candidate intervals that are 50 bp or larger and that do not overlap a chain gap (i.e. have a genotype-phenotype violation) in more than one scorable outgroup chain. In this procedure, a CONDEL can be “called by” (linked to) multiple gene orthologs if they are less than 200 kb from the same candidate region. In Supplementary File 2, we report unique percomorph CONDEL intervals and note all linked orthologs that call a given candidate. The set of pCONDEL-linked orthologs was used for the functional enrichment analysis described in the following section.

#### Functional enrichment test

Using GO Ontology (Ashburner *et al*., 2000; The Gene Ontology Consortium *et al*., 2021) Biological Process Complete annotations (DOI: 10.5281/zenodo.6799722 Released 2022-07-01) and false discovery rate multiple hypothesis testing correction, a PANTHER binomial overrepresentation test (Released 20220712) was performed on the 3,489 `oryLat04` orthologs that yielded one or more pCONDEL candidates in the computational screen (Mi *et al*., 2019). All `oryLat04` protein-coding genes (provided by PANTHER) constituted the background set for the binomial test. See Supplementary File 3 for a complete list of all ontology terms with *q*-value<0.05, all input genes, and all unmapped input genes (which are almost exclusively non-coding RNA genes).

### Phylogenetic tree and branch length calculation

To create a multiple sequence alignment for estimating the phylogenetic distance between species in the screen, we passed the pairwise netted alignments generated by `doBlastzChainNet.pl` into ROAST `v3` (`multiz`) (Blanchette *et al*., 2004). We then used `msa_view` to identify and extract sufficient statistics for four-fold degenerate synonymous sites in coding regions as defined by Ensembl release 98 ASM223467v1 gene models (Cunningham *et al*., 2022). We estimated the rate of neutral evolution at four-fold degenerate sites using `phyloFit` (Hubisz, Pollard and Siepel, 2011) under the REV substitution model, and used `PhyloDM` (Mussig, 2022) to compute pairwise distances between all species in the screen (Supplementary File 4). We based the percomorph phylogenetic tree topology on the consensus of several recent studies (see Fig. 1–fig sup 1 and text sup 1) (Alfaro *et al*., 2018; Hughes *et al*., 2018; Mu *et al*., 2022). `MEGA11 v11.0.13` was used to format and draw the resulting branch length-calibrated tree (Tamura, Stecher and Kumar, 2021).

### Nearest outgroup definition

To identify a given target clade’s nearest outgroup for use in functional experiments, we considered the following factors in the listed order:

1. Proximity based on the most recent common ancestor by phylogenetic tree topology (Fig. 1A)
2. Pairwise genetic distance based on the neutral model of evolution described above (see Fig. 1–Fig. Sup. 1-2, Supplementary File 4)
3. Practical availability of high-quality genomic DNA or adequately preserved tissue samples

### BLAST

`BLASTN v2.7.1+` (task specified to `blastn`) was used to locate regions of sequence similarity between the Japanese Medaka reference genome sequence and the 500 bp and 2.5 kb *PelA* enhancer sequences described by Chan *et al*. (Altschul *et al*., 1997; Chan *et al*., 2010)

### Percomorph tissue and genomic DNA samples

Percomorph genomic DNA was isolated from ethanol-preserved fin, muscle, or liver tissue by incubation in lysis buffer (10 mM Tris, pH 8, 100 mM NaCl, 10 mM EDTA, 0.5% SDS, 500 μg/mL Proteinase K) at 55 °C for 4-12 hours, followed by extraction with 25:24:1 phenol:chloroform:isoamyl alcohol, 1 to 2 washes in chloroform, ethanol precipitation, and resuspension of the DNA pellet in water or TE buffer. Sample and tissue sources are listed in Table S2.

### Species verification by *COI* sequencing

A segment of the mitochondrial gene *cytochrome oxidase subunit I (COI*) was amplified using DreamTaq Green PCR Master Mix (Thermo Scientific catalog # K1081) and primers `percomorph-COI-fwd` and `percomorph-COI-rev` (see Table S5), according to manufacturer recommendations. The subsequent Sanger sequencing reads were cross-referenced with the Barcode of Life Data System (http://v4.boldsystems.org/) to confirm species identity prior to further empirical interrogation of the genetic material (Ratnasingham and Hebert, 2007). *COI* amplicon sequences of *non-Gasterosteus* species have been deposited in GenBank (see Table S2 for accession numbers).

### Sequence verification and cloning of candidate CONDEL regions

We verified the computationally derived pCONDEL candidates near *Pitx1* and *Faf1* by PCR and Sanger sequencing to confirm the DNA sequence represented by the genome assemblies and alignments, according to the following procedure. To compare equivalent genomic sequence between species, we first identified – in Japanese Medaka genomic space – the nearest intervals upstream and downstream of the pCONDEL candidate that exhibit alignment blocks in nearly every outgroup *and* target species with a successful orthologous mapping to *Pitx1* or *Elavl4* and *Faf1*, respectively (see *whole-genome alignment and orthology mapping* methods above). We then used the coordinates of these flanking conservation anchors to extract comparable sequence from each query genome assembly using the script `getConsAnchoredOrthologousFastas.sh` (see code repository).

We designed amplification primers for the extracted assembly sequences using Geneious Prime (Biomatters Ltd) and generated PCR amplicons using Q5 High-Fidelity 2X Master Mix (New England Biolabs catalog # M0492), Phusion High-Fidelity PCR Master Mix with HF Buffer (New England Biolabs catalog # M0531), and/or LongAmp^®^ Hot Start Taq DNA Polymerase with Expand HF Buffer (Roche catalog # 05917131103) according to manufacturer recommendations. The PCR amplicons were then integrated into vector pCR4Blunt-TOPO using the Zero Blunt TOPO PCR Cloning Kit (Invitrogen catalog # 450031) for subsequent Sanger sequencing. Primers introducing homology arms for Gibson Assembly were used to clone candidate sequences into either the GFP reporter vector pT2HE (Howes, Summers and Kingsley, 2017) and/or the luciferase reporter vector pGL4.23[luc2/minP] (Promega, Genbank Accession # DQ904455.1) using NEBuilder HiFi DNA Assembly Master Mix (New England Biolabs catalog # E2621). Reporter constructs and candidate region amplicon sequences have been deposited in GenBank (see Table S2 for accession numbers).

### Threespine Stickleback husbandry

All Threespine Stickleback in this study were raised in 29-gallon tanks under standard aquarium conditions (3.5 g/L Instant Ocean salt, 18°C) and fed live brine shrimp as larvae, then frozen daphnia, bloodworms, and/or mysis shrimp as juveniles and adults – in accordance with the Guide for the Care and Use of Laboratory Animals of the National Institutes of Health under Protocol #13834 of the Institutional Animal Care and Use Committee of Stanford University. All live fish in this study are lab-raised descendants of wild-caught fish. All quantitative *in-vivo* experiments that tested specific hypotheses adhered to ARRIVE guidelines (Percie du Sert *et al*., 2020).

### Enhancer reporter assays in transgenic Threespine Stickleback

Microinjection of freshly fertilized Threespine Stickleback eggs with *Tol2* transposase mRNA and GFP reporter constructs was performed as previously described (Chan *et al*., 2010; Thompson *et al*., 2018). All resulting larvae were raised under standard aquarium conditions to Swarup stage 30 (Swarup, 1958) and then euthanized in 600 mg/L tricaine, pH 7.5 for phenotyping. GFP expression was documented using a Leica MZFLIII fluorescence microscope (Leica Microsystems) fitted with a GFP2 filter and ProgResCF camera (Jenoptik AG). Only fish exhibiting bilateral green eyes (from background expression by the *hsp70* minimal promoter, indicating extent of transgenesis) were phenotyped (Nagayoshi *et al*., 2008). The percomorph *PelA* constructs were assayed on lab-raised pelvic- and caudal-complete stickleback descended from fish collected at Rabbit Slough, Alaska, USA. The construct containing the Japanese Medaka ortholog of the *Faf1* CONDEL candidate region was assayed on lab-raised pelvic- and caudal-complete Threespine Stickleback descended from fish collected at Matadero Creek, California, USA and at Little Campbell River, British Columbia, Canada. The experiment did not include blinding, but post-hoc PCR confirmation of integrated plasmid identity was performed.

### Luciferase enhancer reporter assays

OLHNI-2, a Japanese Medaka fibroblast-like fin-derived cell line, was acquired from RIKEN BioResearch Resource Center (Cell No. RCB2942) and verified by sequencing a segment of the mitochondrial gene *COI* (see *species verification by COI sequencing* above) (Hirayama, Mitani and Watabe, 2006). The cells were cultured using Leibovitz’s L-15 Medium (Gibco Catalog # 11415064) supplemented to 20% fetal bovine serum (ATCC Catalog # 30-2020) and 1x penicillin-streptomycin (Gibco Catalog # 15140122), at 30°C to 33°C and ambient CO_2_ concentration. Firefly and pRL-SV40 renilla luciferase constructs (Promega, Genbank Accession # AF025845) were transfected using Solution SF (Lonza Catalog # V4SC-2096) and program DN-100 of the Amaxa Nucleofector 96-well Shuttle System according to manufacturer recommendations. Each firefly luciferase plasmid was independently amplified, miniprepped (ZymoResearch Catalog # D4210), and tested the following number of times: Marine Stickleback, 12; Yellow Croaker, 33; Green Spotted Puffer, 12; Ocean Sunfish, 8; European Seabass, 10; Gulf Pipefish, 11. Each plasmid preparation replicate was assayed in technical quadruplicate and reported as a single averaged point in Fig. 2D. For inter-experiment normalization, aliquots of a single preparation of the basal firefly vector pGL4.23[luc2/minP] (Promega, Genbank Accession # DQ904455.1) were assayed on each experimental day. Dual-Luciferase Reporter Assay System reagents (Promega Catalog # E1910) and a Dual Injector System for GloMax-Multi Detection System (Promega) luminometer were used to measure luciferase activity. The plasmid locations on each plate were randomized to control for potential positional biases from the multichannel pipettes, nucleofector, and luminometer that were used. Two-tailed Mann Whitney U tests were performed (using python package scipy v1.7.3) to compare the distribution of biological replicates (each the average of technical quadruplicates) between constructs, as this statistical test does not require the data to be normally distributed (Virtanen *et al*., 2020).

### RNAseq

A lab-raised Matadero Creek male Threespine Stickleback and lab-raised Little Campbell River female Threespine Stickleback were crossed to generate one clutch of wildtype F1 hybrid fish. Caudal fin buds were collected into 1.5 mL centrifuge tubes from twelve siblings of the clutch on 10 dpf (after hatching, before lepidotrichia development, Swarup stage 27) and twelve additional siblings on 17 dpf (during lepidotrichia development, Swarup stage 29) (Swarup, 1958). The dissected tissues were immediately placed on dry ice and stored at −80°C until RNA extraction was performed using the RNeasy Plus Micro Kit (Qiagen Catalog # 74034). Each fin bud was first homogenized in 50 μL of Buffer RLT (supplemented with β-mercaptoethanol per kit recommendation) for 1 min at RT in a 1.5 mL centrifuge tube using a cordless motor (VWR Catalog # 47747-370) and pestle (USA Scientific Catalog # 1415-5390). The pestle was rinsed with an additional 300 μL of Buffer RLT into the 1.5mL tube; then the resulting 350 μL volume was drawn twice into a 1 mL luer slip tip tuberculin syringe (BD Catalog # 309659) fitted with a 27G-½” needle. Each RNA sample was eluted in 16 μL of RNAse-free water and quantified using the Qubit RNA High Sensitivity Kit (Invitrogen Catalog # Q32852). A subset of the samples from each developmental stage was quality-checked by Bioanalyzer using the RNA 6000 Pico Kit (Agilent Technologies Catalog # 5067-1513). The resulting RNA integrity numbers were between 8.4 and 10, with most values 9.4 or higher.

Sequencing libraries were generated with the Stranded mRNA Prep kit (Illumina Catalog # 20040532) using 68-80 ng of RNA for 10 dpf samples and 150 ng of RNA for 17 dpf samples. The libraries were sequenced (2 × 150 bp) as a single multiplexed pool on two partial lanes of an Illumina NovaSeq 6000 by Novogene Corporation Inc. The resulting sequencing reads were trimmed with `fastp v0.21.0` (Chen *et al*., 2018) and mapped to Threespine Stickleback genome assembly `gasAcu1-4` via two-pass mapping with STAR v2.7.7a (Dobin *et al*., 2013) according to GATK best practices (Van der Auwera *et al*., 2013). The program `featureCounts v2.0.1` (Liao, Smyth and Shi, 2014) was used to quantify reads mapped per gene based on `gasAcu1` gene annotations (Ensembl release 99) that were lifted to `gasAcu1-4` as previously described (Roberts Kingman *et al*., 2021; Cunningham *et al*., 2022). Each sample’s gene-level read counts were normalized to transcripts per million (TPM) to assess expression levels of genes near *Faf1* at the two developmental stages (Li *et al*., 2010).

### CRISPR-Cas9 genome editing

Single guide RNAs (sgRNAs) for directing Cas9 nuclease activity in Threespine Stickleback were designed using CHOPCHOP (Labun *et al*., 2016, 2019) and the genome assembly `gasAcu1`. Templates for the sgRNAs were synthesized via 2-oligo PCR and used for in-vitro transcription by T7 RNA polymerase, as previously described (Wucherpfennig, Miller and Kingsley, 2019). In addition to sgRNAs targeting candidate regions, an sgRNA for inactivating the *golden* pigment gene *Slc24a5* (Lamason *et al*., 2005) was also included as a visual control for editing status. Table S5 lists all oligonucleotides used to generate sgRNAs.

For recreating the pCONDEL.1189 candidate near *Faf1* in Threespine Stickleback, a 60 bp single-stranded DNA donor template was injected along with the sgRNAs and Cas9 protein. Intended for promoting homology-directed repair, this oligonucleotide was a concatemer of the two 30-mers on either side of the desired 464 bp deletion (see Fig. 4A and Table S5), and the two most 5’ and two most 3’ bases were synthesized to have phosphorothioate (not phosphodiester) bonds to enhance stability *in-vivo* (Integrated DNA Technologies, Inc).

Cas9-2NLS protein (QB3 MacroLab, University of California - Berkeley), sgRNAs, and single-stranded DNA oligonucleotides were injected at the indicated concentrations (Table S3) into freshly fertilized Threespine Stickleback eggs. For experiments in which half of each clutch was injected with reagents to target the pCONDEL.1189 region and the other half was injected with sgRNAs targeting only the *golden* locus, the order of injection was alternated between each of the 17 genome-edited clutches to control for any potentially confounding effects related to the time of injection after fertilization. To verify editing status in the resulting fish, dorsal spine and fin biopsies and/or skin mucus swabs were collected from fish after reaching ≥20 mm standard length and used for subsequent DNA extraction (Breacker *et al*., 2017). All phenotypic scoring was performed at or after fish reached 15 mm standard length; blinding was not performed. All CRISPR-Cas9 editing experiments were performed on either the Matadero Creek and/or Little Campbell River population background. Two-tailed, Boschloo’s Exact tests were performed (using `python` package `scipy` v1.7.3) to ascertain whether the genome editing was significantly associated with phenotypic outcome, as this statistical test is conditioned only on one marginal sum (the number of fish in each treatment or control group) (Boschloo, 1970; Mehrotra, Chan and Berger, 2003; Ludbrook, 2013; Virtanen *et al*., 2020).

#### Sample size calculations

A pilot experiment to target the pCONDEL.1189 candidate region in Threespine Stickleback from Matadero Creek suggested the ectopic caudal ray phenotype occurred in ~5% of injected individuals—and the controls for this pilot experiment consisted of unmodified clutch siblings, which all developed wildtype caudal fins. After completing the pilot experiment, we incidentally observed that a spontaneous caudal fin morphology reminiscent of the pCONDEL.1189-edited phenotype appears at very low frequencies (~0.5%) in wildtype, unmodified Threespine Stickleback from Matadero Creek. Given this finding, we performed power calculations (alpha = 0.05, beta = 0.2, power = 0.8) to determine the appropriate sample size for a replication experiment in which control siblings would also be injected to edit the unrelated *golden* pigment locus to account for possible confounding effects from introducing Cas9 nuclease into the zygote. We report full results of the replication experiment here (involving 17 genome-edited clutches total; see Fig. 4–fig sup 1-2). Because the *golden-edited* controls from the replication experiment above did not suggest confounding effects from the presence of Cas9 nuclease, the subsequent experiment targeting coding exons of *Faf1* used unmodified clutch siblings as controls. A pilot experiment to knockout *Faf1* yielded a similar (~5%) frequency of ectopic caudal ray phenotypes, so we aimed to achieve a similar sample size as above to test the effect of *Faf1* inactivation in a subsequent replication experiment (involving 10 genome-edited clutches total; see Fig. 4–fig sup 3-4).

### Primers and DNA oligonucleotides

Sequences for all DNA oligonucleotides and genotyping primers are listed in Table S5.

### Skeletal preparations

Adult Threespine Stickleback (≥ 30 mm standard length), a Green Spotted Pufferfish, and a juvenile Rice Eel were fixed in 10% neutral buffered formalin for at least 5 days, rinsed twice in distilled water and twice in 30% saturated sodium borate, and then cleared in a solution of 1% porcine trypsin dissolved in 30% saturated sodium borate for 24-48 hours at RT. After equilibrating the cleared specimens to 2% potassium hydroxide (KOH), staining of calcified skeletal tissue was performed by incubation in 0.005% Alizarin Red S dissolved in 2% KOH for 12-24 hours; and then pigment was bleached by incubating the fish in 0.375% KOH, 25% glycerol, and 0.0003% hydrogen peroxide. Juvenile Threespine Stickleback (≤ 20 mm standard length) that required genotyping after skeletal characterization were stained using Walker and Kimmel’s two-color acid-free method (at 100 mM magnesium chloride concentration) (Walker and Kimmel, 2007). Finally, all specimens were passed through a 0.375% KOH:glycerol gradient before imaging and storage in 100% glycerol. An Epson Perfection V800 Photo scanner and a Stemi SV11 stereomicroscope fitted with an AxioCam HRc camera (Carl Zeiss AG), respectively, were used to record brightfield images of adult and juvenile skeletal preparations.

## Code, data, and material availability

A code repository with instructions for replicating the computational screen starting with flat-text genome assembly fasta files is available at https://github.com/bejerano-lab/percomorphCONDELs. To aid users who may not have access to a compute cluster, parts of the screen were implemented to optionally use GNU Parallel (Tange, 2011). Supplementary File 1 lists genome assembly accession identifiers and sources. A copy of pre-computed, genome assembly-derived intermediate inputs (including whole genome alignment chains) for generating the final candidate list has been deposited at https://doi.org/10.5281/zenodo.7140839. Table S2 lists the source of percomorph tissue and genomic DNA used in this study as well as the GenBank accession numbers for the derived DNA sequences and constructs. Raw RNA-seq data are available through National Center for Biotechnology Information (NCBI) BioProject number PRJNA908888. Other materials will be made available upon reasonable request.

## Acknowledgements

We acknowledge the Ohlone and Semiahmoo Nations, from whose traditional territories the Threespine Stickleback populations used in this study were derived. We are grateful to the following people for providing non-stickleback tissue and/or DNA samples used in this study: Andrew Bentley, Pierre-Alexandre Gagnaire, François Bonhomme, Jessica Muniz, Angel Seery, Leo Nico, Chris Smith, Scott Collenberg, Carol Cozzi-Schmarr, Xuegeng Wang, Ramji Bhandari, Susan L. Bassham, William A. Cresko, Molly Schumer, Shreya Banerjee, Anne Brunet, Rahul Nagvekar, Leonor Cancela, and Vincent Laizé. We thank the numerous researchers who generated and made freely available the genome assemblies used in this study; the UCSC Genome Browser Team and Hiram Clawson for the generously shared wisdom and tools they have developed for visualizing and analyzing genomic data; Dave Catania for assistance in obtaining radiographs of various fish species; Julia Wucherpfennig, Abbey Thompson, Ken Thompson, and Dolph Schluter for help collecting sticklebacks from Little Campbell River; Sarah Frail for contributions to initial CRISPR/Cas9 experiments; William Talbot for generously lending imaging equipment; the team of fish feeders for their husbandry support; and all members of the Bejerano and Kingsley Labs (especially Mark Berger, Amir Marcovitz, Johannes Birgmeier, Karthik Jagadeesh, Whitney Heavner, Boyoung Yoo, Yosuke Tanigawa, Janet Song, Amy Herbert, and Julia Wucherpfennig) as well as Michael A. Bell, William Talbot, and Margaret Fuller for their helpful suggestions. This work was supported by a National Science Foundation Graduate Research Fellowship (1656518), NIH training grant (5T32GM007790), and Stanford CEHG Fellowship to H.I.C.; D.M.K. is an investigator of the Howard Hughes Medical Institute.

## Author Contributions

H.I.C., G.B., and D.M.K. conceptualized the computational screen. Y.T. provided code for identifying orthologous alignment chains. H.I.C. curated and aligned the publicly available genome assemblies and wrote the code for performing the computational screen. H.I.C. and D.M.K. conceptualized the functional experiments for testing the candidate regions. H.I.C. carried out the investigations and performed all data analysis and visualization. H.I.C. and D.M.K. wrote the manuscript with input from all authors. G.B. and D.M.K. supervised the study, managed resources, and acquired funding.

