## Supplemental Figures, Text, and Tables for "Whole-genome comparisons identify repeated regulatory changes underlying convergent appendage evolution in diverse fish lineages"

#### Figure 1–figure supplement 1

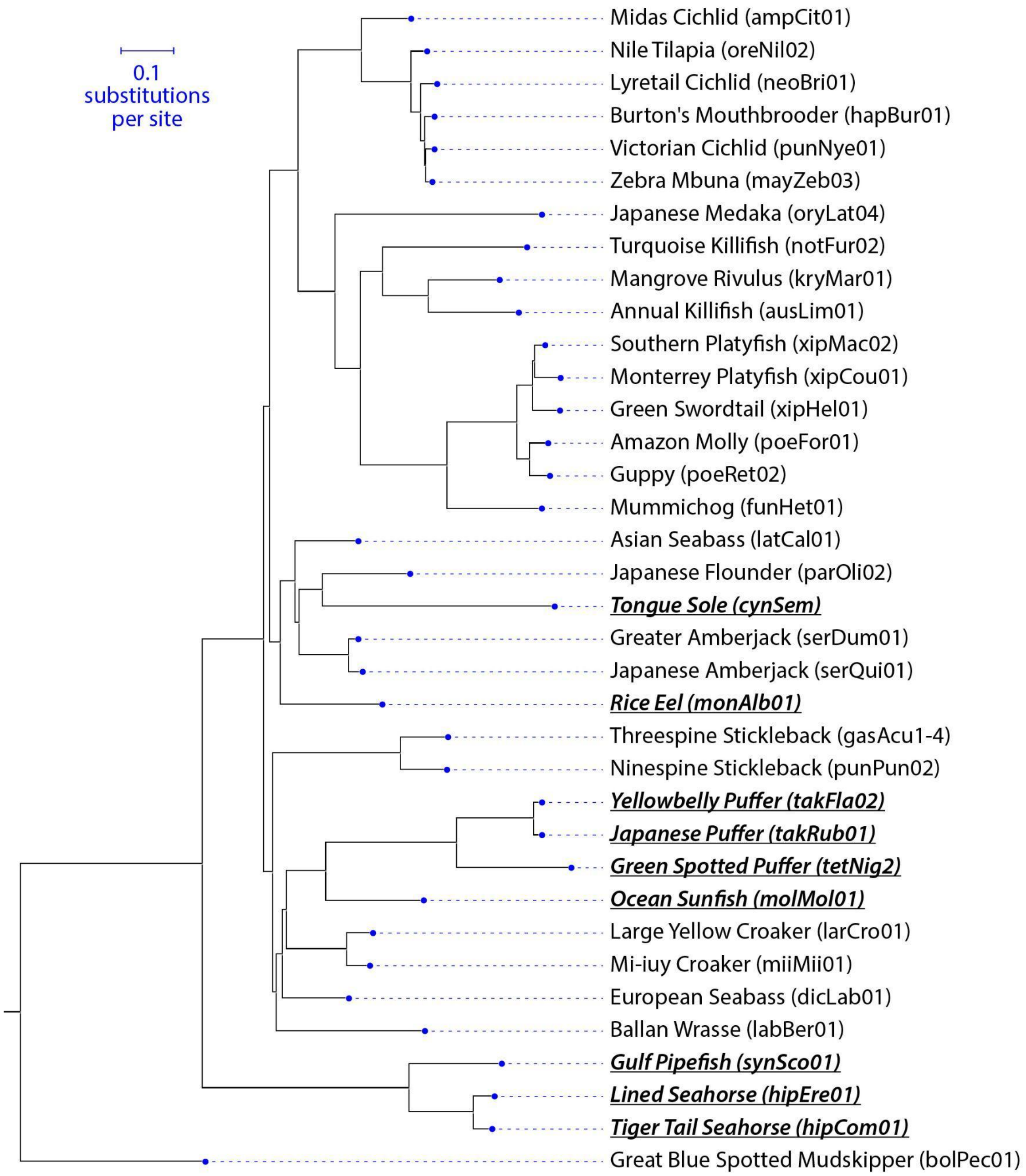


**Figure 1–figure supplement 1. Branch length-calibrated tree.** Common names (with associated genome assembly abbreviations) for all 36 species in the computational screen are listed (also see Supplemental File 1). Target species are underlined and italicized. The topology of this tree is based on a consensus of several recent studies (Alfaro *et al.*, 2018; Hughes *et al.*, 2018; Mu *et al.*, 2022).

### Figure 1–text supplement 1

**Figure 1–text supplement 1. Branch lengths for trees shown in Fig. 1A and Fig. 1–fig sup 1**

```
ALPHABET: A C G T
ORDER: 0
SUBST_MOD: REV
BACKGROUND: 0.190773 0.360458 0.235183 0.213586
RATE_MAT:
  -1.168358    0.296539    0.622435    0.249384
   0.156944   -0.785009    0.199225    0.428841
   0.504898    0.305345   -1.025896    0.215653
   0.222747    0.723732    0.237459   -1.183938
TREE:
((((((ampCit01:0.0887886,(oreNil02:0.0232145,(neoBri01:0.0255143,(hapBur01:0.0112809,(punNye01:0.0105135,mayZe
b03:0.00683988):0.00242368):0.00857014):0.0188007):0.0940882):0.119616,(oryLat04:0.385762,((notFur02:0.267095,
(ausLim01:0.166114,kryMar01:0.130003):0.0857237):0.042407,((((xipMac02:0.0143759,xipCou01:0.0434946):0.0039567
9,xipHel01:0.0468436):0.0309337,(poeFor01:0.0306962,poeRet02:0.0337162):0.0245541):0.131882,funHet01:0.174194)
:0.164058):0.0487246):0.0707918):0.0535661,((latCal01:0.114761,((parOli02:0.160689,cynSem:0.433672):0.0437825,
(serQui01:0.0202901,serDum01:0.0129067):0.0946104):0.00742124):0.0278147,monAlb01:0.188107):0.0202366):0.01178
74,((gasAcu14:0.0832039,punPun02:0.0819135):0.241185,((((((takFla02:0.00916032,takRub01:0.0091555):0.146708,te
tNig2:0.212735):0.248094,molMol01:0.180985):0.0734705,(larCro01:0.0435642,miiMii01:0.0373942):0.114976):0.0082
0025,dicLab01:0.119023):0.0125103,labBer01:0.276969):0.00632666):0.0188741):0.116492,(synSco01:0.170947,(hipEr
e01:0.0331536,hipCom01:0.0298633):0.12246):0.392849):0.343345,bolPec01:0.343345);
```

#### Figure 1–figure supplement 2

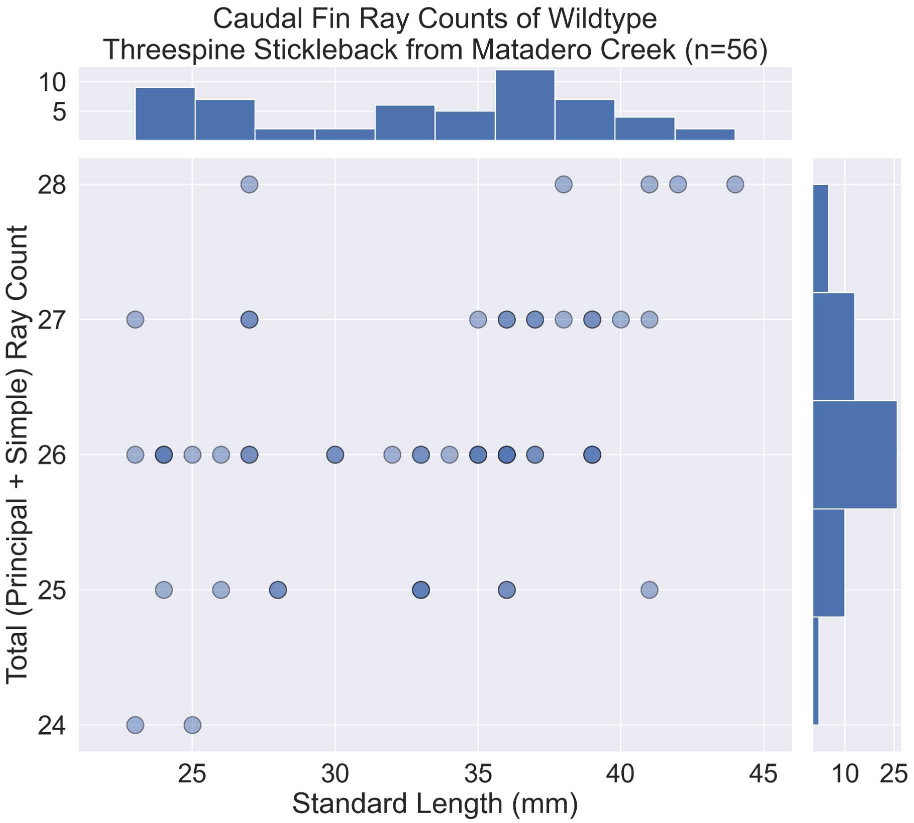


**Figure 1–figure supplement 2. Caudal ray counts in wildtype Matadero Creek Threespine Stickleback.** Total *segmented principal* and *unsegmented simple* caudal ray (lepidotrichia) counts plotted against fish standard length. Data are from n=56 lab-reared wildtype Threespine Stickleback descendants of fish collected from Matadero Creek, California.

#### Figure 4–figure supplement 1

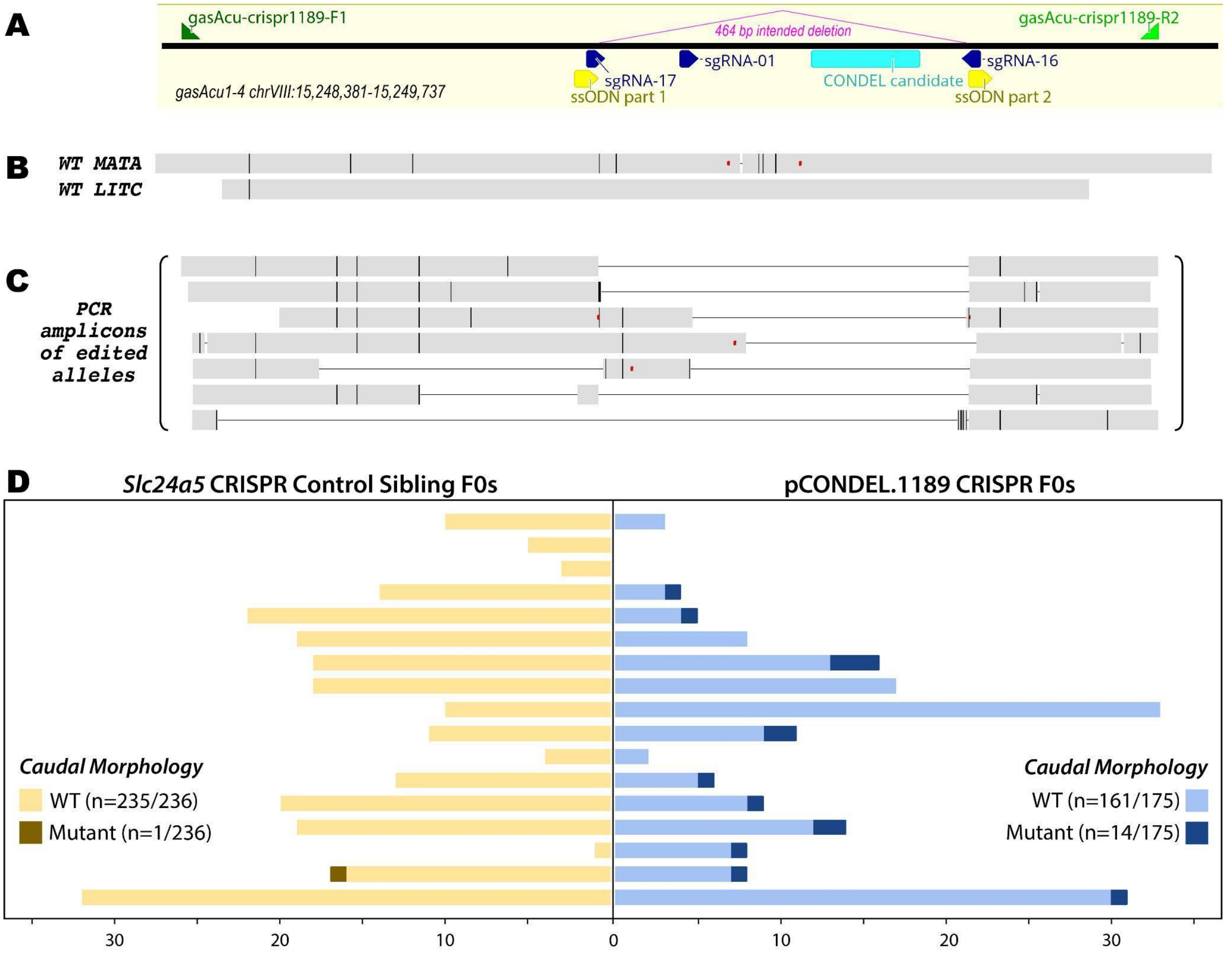


**Figure 4–figure supplement 1. CRISPR-Cas9 deletions and morphological effects of recapitulating the pCONDEL.1189 evolutionary lesion. A**, Schematic of the CRISPR-targeted pCONDEL.1189 genomic region in Threespine Stickleback (`gasAcu1-4`) genomic space. PCR primers are shown as green triangles, sgRNAs are shown as dark blue triangles, and the injected 60 bp single-stranded DNA oligonucleotide is a concatemer of the two yellow triangles (Table S5). **B**, Sanger-sequenced amplicons from wildtype (WT) descendants of Matadero Creek, California (MATA) and Little Campbell River, Canada (LITC). **C**, Examples of Sanger-sequenced amplicons recovered from crispants injected with Cas9 protein and the reagents illustrated above (see Table S3 for injected concentrations and Table S5 for oligonucleotide sequences). In **B** and **C**, black vertical lines indicate nucleotide substitutions relative to the `gasAcu1-4` reference sequence; horizontal black lines represent deleted sequence; red dots denote insertions; and light gray bars illustrate reference-identical sequence. **D**, Paired bar graphs showing the number of scored *Slc24a5* CRISPR control siblings on the left, and pCONDEL.1189 CRISPR F0s on the right for 17 clutches (each row represents one clutch). Darker and lighter shaded portions of the bar graphs, respectively, indicate the number of individuals with ectopic dorsal segmented rays (as in Fig. 4C) or with wildtype caudal morphology (as in Fig. 4B).

### Figure 4–figure supplement 2

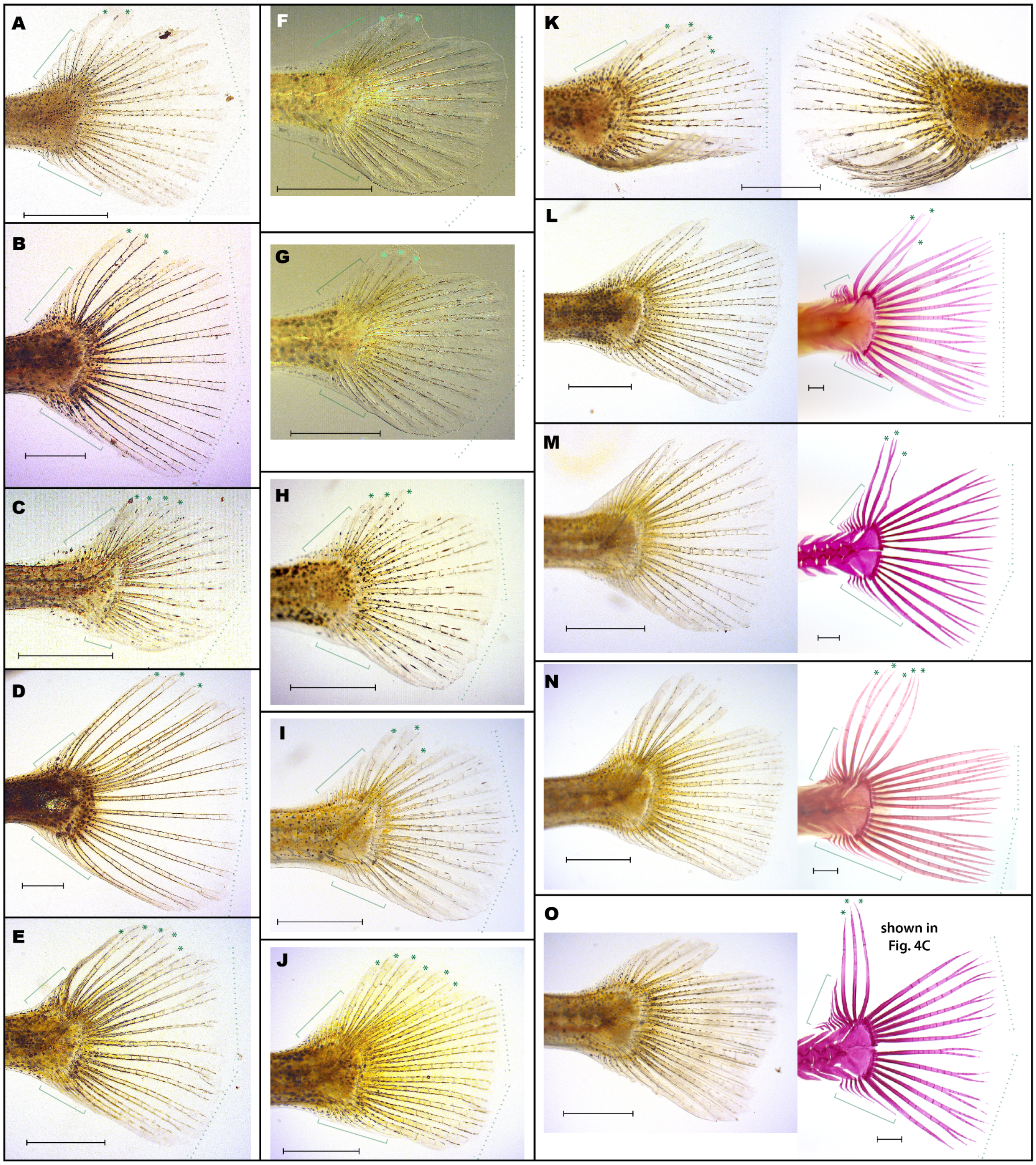


**Figure 4–figure supplement 2. Fish from pCONDEL.1189-targeting experiments with ectopic dorsal rays. A**, The single *Slc24a5*-targeted control sibling (out of 236 total) that developed ectopic dorsal caudal rays. **B-O**, All 14 enhancer deletion F0 crispant fish (out of 175 total) reported in Fig. 4–fig sup 1D to have ectopic dorsal rays. Panel **K** shows the left and right sides of one fish with a curled tail fin. **L-O** each shows two photos of a single fish – at an earlier and later developmental stage from left to right, respectively. Dashed brackets denote typical, segmented *principal* rays on the dorsal and ventral halves of the tail fin; solid brackets denote unsegmented *simple* rays; asterisks (*) denote ectopic segmented rays. All scale bars are 1 mm.

#### Figure 4–figure supplement 3

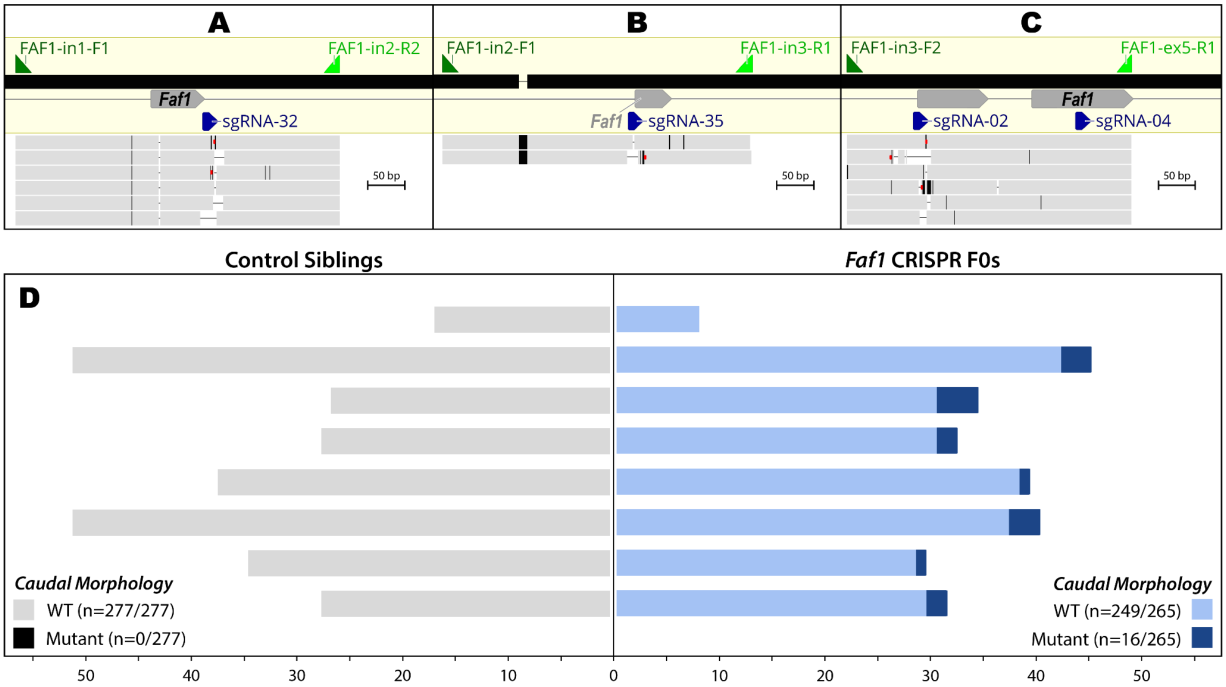


**Figure 4–figure supplement 3. CRISPR-Cas9 lesions and morphological effects of disrupting *Faf1* coding regions. A-C**, Schematics of exons 2-5 of *Faf1* in Threespine Stickleback (`gasAcu1-4`) sequence space. PCR primers are shown as green triangles. Examples of PCR amplicons from fish injected with Cas9 protein and all 4 of the sgRNAs (dark blue triangles) illustrated (see Table S3 for injected concentrations and Table S5 for oligonucleotide sequences). Black vertical lines indicate nucleotide substitutions relative to the `gasAcu1-4` reference sequence; horizontal black lines represent deleted/missing sequence; red dots denote insertions; and light gray bars illustrate reference-identical sequence. Numerous editing events were detected at the target of sgRNA-32 in **A** and sgRNA-02 in **C**. Very few editing events were detected at the target of sgRNA-35 in **B**, and no conclusive editing events were detected at the target of sgRNA-04 in **C**. **D**, Paired bar graphs showing the number of scored control siblings on the left, and *Faf1*-targeted siblings on the right, from 10 clutches. Two sets of smaller clutches were combined for practical husbandry purposes, so 8 total rows are shown. Darker and lighter shaded portions of the bar graphs, respectively, indicate the number of individuals with ectopic dorsal segmented rays (as in Fig. 4F) or with wildtype caudal morphology (as in Fig. 4E).

#### Figure 4–figure supplement 4

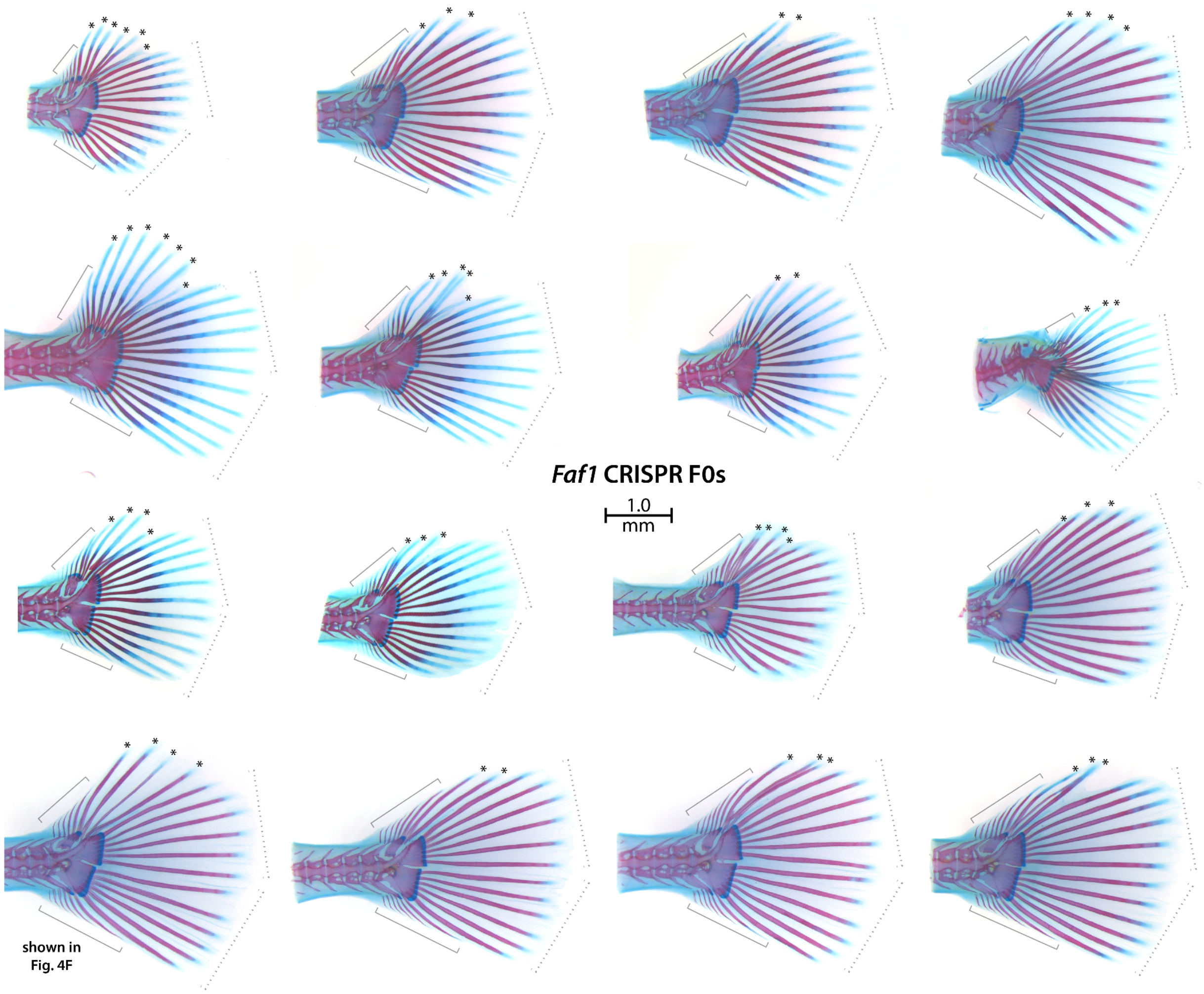


**Figure 4–figure supplement 4. Fish from *Faf1*-targeting experiments with ectopic dorsal rays.** All 16 *Faf1*-targeted F0 crispant fish reported in Fig. 4–fig sup 3D to have ectopic dorsal rays. Dashed brackets denote typical, segmented *principal* rays on the dorsal and ventral halves of the tail fin; solid brackets denote unsegmented *simple* rays; asterisks (*) denote ectopic segmented rays.

#### Figure 4–figure supplement 5

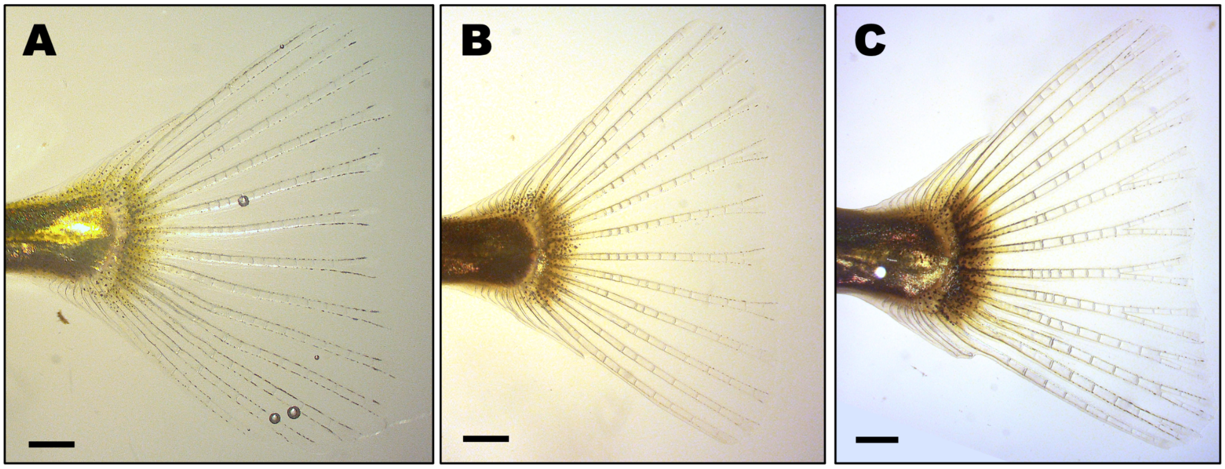


**Figure 4–figure supplement 5. Wildtype caudal morphology in fish with stably transmitted pCONDEL.1189 deletion alleles.** Mosaic F0 CRISPR founders were intercrossed to generate F1 fish carrying different combinations of wildtype and pCONDEL.1189 deletion alleles. Representative fish from three clutches that all had wildtype caudal morphology, including individuals with two intact pCONDEL.1189 alleles (**A**), fish homozygous for a 927 bp pCONDEL.1189 deletion allele (**B**), and compound heterozygotes carrying 927bp and 229bp pCONDEL.1189 deletion alleles (**C**). All deletion alleles in the fish shown in **B** and **C** remove the entire 135 bp pCONDEL.1189 sequence. All scale bars are 1 mm.

### Supplemental Tables

**Table S1:** References for pelvic and caudal fin status of key target and outgroup species.

| Scientific name | Common name | # of pelvic fins | Total # of caudal fin rays | Reference |
|---|---|---|---|---|
| *Cynoglossus crepida* | Congener of Tongue Sole | 1 | 8 | [1] |
| *Cynoglossus semilaevis* | Tongue Sole | 1 | 10 | [2] |
| *Cynoglossus* sp. | Congener of Tongue Sole | 1 | 8-12 | [3] |
| *Cynoglossus westraliensis* | Congener of Tongue Sole | 1 | 8 | [4] |
| *Cynoglossus yokomaru* | Congener of Tongue Sole | 1 | 10 | [5] |
| *Dicentrarchus labrax* | European Seabass | 2 | 16-18 principal + numerous simple rays | [6] |
| *Dicentrarchus labrax* | European Seabass | [not mentioned] | 17 principal, 29-31 simple | [7, 8] |
| *Gasterosteus aculeatus* | Threespine Stickleback (Matadero Creek) | 2 | $\geq$24 | This study: see Fig 1–Sup. Fig.2 |
| *Gasterosteus aculeatus* | Threespine Stickleback | [not mentioned] | 22-31 for fish $\geq$15mm | [9] |
| *Gasterosteus aculeatus* | Threespine Stickleback | [not mentioned] | 24 | [10] |
| *Hippocampus comes* | Tiger Tail Seahorse | 0 | [not mentioned] | [11] |
| *Hippocampus erectus* | Lined Seahorse | 0 | [not mentioned] | [11] |
| *Hippocampus* sp. | Congener of seahorses | 0 | 0-3 | [12] |
| *Larimichthys crocea* | Yellow Croaker | 2 | 35 | [13] |
| *Mola mola* | Ocean Sunfish | 0 | 0 | [11, 14, 15] |
| *Monopterus albus* | Rice Eel | 0 | 0 | [16, 17] |
| *Oryzias latipes* | Japanese Medaka | [not mentioned] | 20-24 | [18] |
| *Paralichthys olivaceus* | Japanese Flounder | 2 | 18-19 | [19] |
| *Paralichthys olivaceus* | Japanese Flounder | 2 | [not mentioned] | [20] |
| *Paralichthys* sp. | Congener of Japanese Flounder | 2 | 18 | [21] |
| *Seriola dumerili* | Greater Amberjack | 2 | 39 | [22] |
| *Syngnathus scovelli* | Gulf Pipefish | 0 | [not mentioned] | [11] |
| *Syngnathus schlegeli* | Congener of Gulf Pipefish | [not mentioned] | 9-10 | [23] |
| Tetraodontoidea | Superfamily of pufferfish | 0 | 9-11 | [14] |
| Tetraodontidae | Family of pufferfish | 0 | 10 | [11] |
| Tetraodontidae | Family of pufferfish | [not mentioned] | 11 | [8] |

**Table S2: Tissue, cell line, and genomic DNA sources.** GenBank accession numbers for DNA sequences and constructs derived from each source are shown, grouped by the indicated genomic locus from which they were amplified. *COI*, cytochrome oxidase subunit 1.

| | | | | GENBANK ACCESSION NUMBERS | | |
|---|---|---|---|---|---|---|
| **Species** | **Common Name** | **Item and Source** | **Voucher ID** | ***COI*** | **pCONDEL.329** | **pCONDEL.1189** |
| *Dicentrarchus labrax* | European Seabass | Tissue and purified genomic DNA gift from P. Gagnaire and F. Bonhomme (University of Montpellier) | n/a | OP896983<br>OP896984 | OP934176 | OP934193<br>OP934194 |
| *Gasterosteus aculeatus* | Threespine Stickleback (Little Campbell River) | [lab-raised, this study] | n/a | n/a | n/a | OP934188 |
| *Gasterosteus aculeatus* | Threespine Stickleback (Matadero Creek) | [lab-raised, this study] | n/a | n/a | n/a | OP934187 |
| *Gasterosteus aculeatus* | Threespine Stickleback (Salmon River, Rabbit Slough) | Salmon River BAC clone GU130435<br>[lab-raised, this study] | n/a | n/a | OP934180<br>OP934181 | n/a |
| *Hippocampus erectus* | Lined Seahorse | Tissue gift from J. Muniz, A. Seery, & C. Cozzi-Schmarr (Ocean Rider Seahorse Farm, Hawaii) | n/a | OP896990 | OP934183<br>OP934184 | OP934185<br>OP934186 |
| *Larimichthys crocea* | Large Yellow Croaker | Purchased from 99 Ranch Market in 2019 | n/a | OP896988 | OP934177<br>OP934178 | OP934189 |
| *Mola mola* | Ocean Sunfish | Tissue gift from the Ichthyology Collection, University of Kansas, Biodiversity Institute (KUBI Ichthyology) | KU2979 | MK144137 | OP934174 | OP934191 |
| *Monopterus albus* | Rice Eel | Tissue gift from L. Nico, C. Smith, & S. Collenberg (US Geological Survey) | LGN12-10 | OP896985 | n/a | OP934192 |
| *Oryzias latipes* | Japanese Medaka (Hd-rR strain) | Tissue and genomic DNA gift from X. Wang and R. Bhandari (UNC Greensboro) | n/a | OP896987 | n/a | OP934173<br>OP934190 |
| *Oryzias latipes* | Japanese Medaka OLHNI-2 cell line | RIKEN BioResearch Resource Center (Japan) | RCB2942 | OP896986 | n/a | n/a |
| *Seriola dumerili* | Greater Amberjack | Tissue gift from the Ichthyology Collection, University of Kansas, Biodiversity Institute (KUBI Ichthyology) | KU5141<br>KU5170 | MK144139<br>MK144135 | n/a | OP934195 |
| *Syngnathus scovelli* | Gulf Pipefish | Genomic DNA gift from S. L. Bassham and W. A. Cresko (University of Oregon) | n/a | OP896989 | OP934182 | n/a |
| *Takifugu rubripes* | Japanese Puffer | Tissue gift from the Ichthyology Collection, University of Kansas, Biodiversity Institute (KUBI Ichthyology) | KU3490 | MK144136 | OP934200 | OP934201 |
| *Tetraodon nigroviridis* | Green Spotted Puffer | Purchased from AZ Aquatic Gardens [24] in 2019 | n/a | OP896980<br>OP896981<br>OP896982 | OP934175<br>OP934179<br>OP934198<br>OP934199 | OP934196<br>OP934197 |

**Table S3:** CRISPR/Cas9 injection mixes.

| Injection Group | Component | Final concentration in injection mix (ng/$\mu$L) | Associated Oligo ID (see Table S5) |
|---|---|---|---|
| Delete CONDEL region near *Faf1* | Cas9-2NLS | 950.00 | n/a |
| | Faf1CONDEL ssDNA | 20.00 | Faf1CONDEL_ssODN |
| | FAF1-01 sgRNA | 43.75 | sgRNA-01 |
| | FAF1-16 sgRNA | 150.00 | sgRNA-16 |
| | FAF1-17 sgRNA | 150.00 | sgRNA-17 |
| | *Slc24a5* sgRNA | 34.15 | sgRNA-SLC24A5 |
| *Slc24a5* CRISPR knockout control | Cas9-2NLS | 950.00 | n/a |
| | Faf1CONDEL ssDNA | 20.00 | Faf1CONDEL_ssODN |
| | *Slc24a5* sgRNA | 377.90 | sgRNA-SLC24A5 |
| *Faf1* CRISPR knockout | Cas9-2NLS | 937.50 | n/a |
| | *Slc24a5* sgRNA | 40.00 | sgRNA-SLC24A5 |
| | FAF1-02 sgRNA | 65.00 | sgRNA-02 |
| | FAF1-04 sgRNA | 65.00 | sgRNA-04 |
| | FAF1-32 sgRNA | 65.00 | sgRNA-32 |
| | FAF1-35 sgRNA | 65.00 | sgRNA-35 |

**Table S4:** Summary statistics for sliding windows used to compute conserved regions.

| Genome assembly aligned to oryLat04 | Siding window size (10, 25, 50, 100 bp) | Number of tied windows added | Percent identity of least-conserved window included | Total number of windows included after adding ties | Coverage of oryLat04 after including ties |
|---|---|---|---|---|---|
| ampCit01 | 10 | 14,072,787 | 90% | 32,138,846 | 7.79% |
| ampCit01 | 25 | 3,865,919 | 88% | 21,312,301 | 5.82% |
| ampCit01 | 50 | 62,054 | 86% | 17,108,604 | 5.01% |
| ampCit01 | 100 | 857,808 | 81% | 15,577,367 | 5.23% |
| ausLim01 | 10 | 3,948,786 | 90% | 23,324,463 | 5.82% |
| ausLim01 | 25 | 1,558,266 | 84% | 20,978,969 | 5.29% |
| ausLim01 | 50 | 705,663 | 80% | 19,704,764 | 5.14% |
| ausLim01 | 100 | 334,819 | 74% | 16,098,009 | 5.09% |
| bolPec01 | 10 | 5,352,533 | 80% | 25,347,326 | 6.06% |
| bolPec01 | 25 | 3,346,874 | 72% | 24,963,848 | 5.56% |
| bolPec01 | 50 | 503,793 | 70% | 20,801,510 | 5.09% |
| bolPec01 | 100 | 222,797 | 64% | 16,522,092 | 5.06% |
| cynSem | 10 | 650,574 | 90% | 19,315,715 | 5.15% |
| cynSem | 25 | 2,988,410 | 80% | 22,635,657 | 5.52% |
| cynSem | 50 | 1,914,327 | 76% | 20,925,087 | 5.36% |
| cynSem | 100 | 490,321 | 71% | 16,080,465 | 5.12% |
| dicLab01 | 10 | 21,153,704 | 90% | 37,649,954 | 9.56% |
| dicLab01 | 25 | 7,348,604 | 88% | 23,962,217 | 6.59% |
| dicLab01 | 50 | 2,764,991 | 86% | 19,201,504 | 5.62% |
| dicLab01 | 100 | 568,002 | 83% | 14,926,303 | 5.15% |
| funHet01 | 10 | 4,479,023 | 90% | 24,243,329 | 5.89% |
| funHet01 | 25 | 3,239,871 | 84% | 22,383,324 | 5.58% |
| funHet01 | 50 | 2,441,583 | 80% | 20,867,043 | 5.47% |
| funHet01 | 100 | 177,199 | 75% | 15,450,144 | 5.04% |
| gasAcu14 | 10 | 8,173,185 | 90% | 26,119,917 | 6.82% |
| gasAcu14 | 25 | 4,886,414 | 84% | 22,995,029 | 5.95% |
| gasAcu14 | 50 | 786,697 | 82% | 18,610,779 | 5.16% |
| gasAcu14 | 100 | 572,246 | 77% | 15,552,985 | 5.15% |
| hapBur01 | 10 | 14,696,278 | 90% | 32,358,250 | 8.03% |
| hapBur01 | 25 | 3,407,200 | 88% | 20,825,668 | 5.74% |
| hapBur01 | 50 | 3,374,405 | 84% | 20,457,531 | 5.71% |
| hapBur01 | 100 | 677,713 | 81% | 15,417,199 | 5.18% |
| hipCom01 | 10 | 6,546,452 | 80% | 26,606,146 | 6.26% |
| hipCom01 | 25 | 468,825 | 76% | 21,971,049 | 5.08% |
| hipCom01 | 50 | 1,187,502 | 70% | 21,456,403 | 5.23% |
| hipCom01 | 100 | 135,044 | 64% | 16,262,575 | 5.03% |
| hipEre01 | 10 | 6,327,045 | 80% | 26,389,929 | 6.22% |
| hipEre01 | 25 | 247,479 | 76% | 21,777,657 | 5.04% |
| hipEre01 | 50 | 1,004,461 | 70% | 21,308,876 | 5.19% |
| hipEre01 | 100 | 70,451 | 64% | 16,231,922 | 5.02% |
| kryMar01 | 10 | 14,311,684 | 90% | 32,002,824 | 7.96% |
| kryMar01 | 25 | 3,046,598 | 88% | 20,524,531 | 5.66% |
| kryMar01 | 50 | 2,856,175 | 84% | 20,041,941 | 5.60% |

*Continued on next page*

**Table S4 – continued from previous page**

| Genome assembly aligned to oryLat04 | Siding window size (10, 25, 50, 100 bp) | Number of tied windows added | Percent identity of least-conserved window included | Total number of windows included after adding ties | Coverage of oryLat04 after including ties |
|---|---|---|---|---|---|
| kryMar01 | 100 | 187,279 | 81% | 14,940,164 | 5.05% |
| labBer01 | 10 | 12,756,378 | 90% | 30,217,543 | 7.82% |
| labBer01 | 25 | 1,121,471 | 88% | 18,518,923 | 5.25% |
| labBer01 | 50 | 777,279 | 84% | 17,871,905 | 5.17% |
| labBer01 | 100 | 824,486 | 79% | 15,439,202 | 5.22% |
| larCro01 | 10 | 19,454,300 | 90% | 37,042,241 | 9.02% |
| larCro01 | 25 | 5,681,994 | 88% | 23,386,277 | 6.18% |
| larCro01 | 50 | 1,106,193 | 86% | 18,529,470 | 5.24% |
| larCro01 | 100 | 452,433 | 82% | 15,557,922 | 5.12% |
| latCal01 | 10 | 23,416,069 | 90% | 39,658,644 | 10.04% |
| latCal01 | 25 | 8,861,270 | 88% | 25,247,455 | 6.91% |
| latCal01 | 50 | 3,960,376 | 86% | 20,172,309 | 5.89% |
| latCal01 | 100 | 1,297,916 | 83% | 15,485,868 | 5.36% |
| mayZeb03 | 10 | 13,959,746 | 90% | 32,046,608 | 7.77% |
| mayZeb03 | 25 | 3,723,013 | 88% | 21,152,282 | 5.80% |
| mayZeb03 | 50 | 3,730,599 | 84% | 20,708,251 | 5.78% |
| mayZeb03 | 100 | 685,940 | 81% | 15,301,383 | 5.18% |
| miiMii01 | 10 | 15,636,572 | 90% | 32,965,956 | 8.36% |
| miiMii01 | 25 | 3,472,642 | 88% | 20,930,878 | 5.75% |
| miiMii01 | 50 | 3,289,211 | 84% | 20,497,835 | 5.69% |
| miiMii01 | 100 | 552,935 | 81% | 15,488,299 | 5.15% |
| molMol01 | 10 | 10,315,861 | 90% | 27,810,890 | 7.32% |
| molMol01 | 25 | 6,986,555 | 84% | 24,502,017 | 6.36% |
| molMol01 | 50 | 2,615,683 | 82% | 19,976,627 | 5.54% |
| molMol01 | 100 | 27,167 | 79% | 14,803,878 | 5.01% |
| monAlb01 | 10 | 13,932,349 | 90% | 30,979,175 | 8.12% |
| monAlb01 | 25 | 1,822,691 | 88% | 18,872,747 | 5.41% |
| monAlb01 | 50 | 1,778,630 | 84% | 18,703,966 | 5.38% |
| monAlb01 | 100 | 1,123,912 | 80% | 15,741,178 | 5.30% |
| neoBri01 | 10 | 12,229,443 | 90% | 30,618,068 | 7.42% |
| neoBri01 | 25 | 2,537,938 | 88% | 20,232,883 | 5.54% |
| neoBri01 | 50 | 2,519,111 | 84% | 19,792,998 | 5.53% |
| neoBri01 | 100 | 907,994 | 80% | 15,700,664 | 5.24% |
| notFur02 | 10 | 2,975,653 | 90% | 22,970,278 | 5.59% |
| notFur02 | 25 | 1,826,110 | 84% | 21,390,479 | 5.33% |
| notFur02 | 50 | 1,420,810 | 80% | 20,300,027 | 5.27% |
| notFur02 | 100 | 44,489 | 75% | 15,698,318 | 5.01% |
| oreNil02 | 10 | 15,361,710 | 90% | 33,191,675 | 8.05% |
| oreNil02 | 25 | 4,676,831 | 88% | 21,924,498 | 6.00% |
| oreNil02 | 50 | 663,857 | 86% | 17,516,078 | 5.15% |
| oreNil02 | 100 | 60,859 | 82% | 14,596,238 | 5.02% |
| parOli02 | 10 | 15,424,400 | 90% | 32,547,196 | 8.38% |
| parOli02 | 25 | 3,104,008 | 88% | 20,249,671 | 5.69% |
| parOli02 | 50 | 2,898,380 | 84% | 19,748,597 | 5.62% |

*Continued on next page*

Table S4 – continued from previous page

| Genome assembly aligned to oryLat04 | Siding window size (10, 25, 50, 100 bp) | Number of tied windows added | Percent identity of least-conserved window included | Total number of windows included after adding ties | Coverage of oryLat04 after including ties |
|---|---|---|---|---|---|
| parOli02 | 100 | 257,238 | 81% | 14,742,233 | 5.07% |
| poeFor01 | 10 | 7,691,039 | 90% | 26,893,407 | 6.52% |
| poeFor01 | 25 | 6,474,480 | 84% | 24,749,061 | 6.17% |
| poeFor01 | 50 | 2,337,774 | 82% | 20,025,703 | 5.47% |
| poeFor01 | 100 | 682,511 | 77% | 15,492,781 | 5.18% |
| poeRet02 | 10 | 7,659,353 | 90% | 26,759,228 | 6.52% |
| poeRet02 | 25 | 6,443,158 | 84% | 24,604,838 | 6.18% |
| poeRet02 | 50 | 2,307,940 | 82% | 19,899,431 | 5.47% |
| poeRet02 | 100 | 697,501 | 77% | 15,410,228 | 5.18% |
| punNye01 | 10 | 13,197,344 | 90% | 31,399,819 | 7.62% |
| punNye01 | 25 | 3,203,697 | 88% | 20,721,252 | 5.69% |
| punNye01 | 50 | 3,214,249 | 84% | 20,281,700 | 5.67% |
| punNye01 | 100 | 309,951 | 81% | 14,971,955 | 5.08% |
| punPun02 | 10 | 9,095,897 | 90% | 26,997,653 | 7.00% |
| punPun02 | 25 | 5,724,424 | 84% | 23,707,510 | 6.11% |
| punPun02 | 50 | 1,404,986 | 82% | 19,079,773 | 5.29% |
| punPun02 | 100 | 848,200 | 77% | 15,742,006 | 5.22% |
| serDum01 | 10 | 22,996,454 | 90% | 39,270,774 | 9.95% |
| serDum01 | 25 | 8,685,504 | 88% | 25,087,116 | 6.86% |
| serDum01 | 50 | 3,888,105 | 86% | 20,149,297 | 5.87% |
| serDum01 | 100 | 1,385,341 | 83% | 15,602,792 | 5.38% |
| serQui01 | 10 | 22,366,299 | 90% | 38,715,761 | 9.82% |
| serQui01 | 25 | 8,213,483 | 88% | 24,701,450 | 6.77% |
| serQui01 | 50 | 3,505,604 | 86% | 19,827,653 | 5.78% |
| serQui01 | 100 | 1,084,960 | 83% | 15,360,789 | 5.30% |
| synSco01 | 10 | 4,981,426 | 80% | 25,381,300 | 5.95% |
| synSco01 | 25 | 3,300,615 | 72% | 25,061,917 | 5.55% |
| synSco01 | 50 | 247,749 | 70% | 20,706,083 | 5.05% |
| synSco01 | 100 | 350,352 | 63% | 16,681,006 | 5.09% |
| takFla02 | 10 | 1,974,966 | 90% | 20,678,470 | 5.44% |
| takFla02 | 25 | 5,311,441 | 80% | 24,562,759 | 5.93% |
| takFla02 | 50 | 1,762,231 | 78% | 20,472,896 | 5.34% |
| takFla02 | 100 | 448,663 | 74% | 15,921,427 | 5.11% |
| takRub01 | 10 | 1,955,663 | 90% | 20,692,350 | 5.44% |
| takRub01 | 25 | 5,262,609 | 80% | 24,562,134 | 5.92% |
| takRub01 | 50 | 1,681,694 | 78% | 20,452,214 | 5.32% |
| takRub01 | 100 | 372,479 | 74% | 15,896,061 | 5.10% |
| tetNig2 | 10 | 11,477,305 | 80% | 31,009,353 | 7.17% |
| tetNig2 | 25 | 234,709 | 80% | 20,864,475 | 5.04% |
| tetNig2 | 50 | 1,931,352 | 74% | 21,681,785 | 5.37% |
| tetNig2 | 100 | 253,266 | 70% | 16,229,452 | 5.06% |
| xipCou01 | 10 | 119,746 | 90% | 20,560,379 | 5.02% |
| xipCou01 | 25 | 3,824,517 | 80% | 24,277,168 | 5.62% |
| xipCou01 | 50 | 402,531 | 78% | 19,863,059 | 5.08% |

*Continued on next page*

Table S4 – continued from previous page

| Genome assembly aligned to oryLat04 | Siding window size (10, 25, 50, 100 bp) | Number of tied windows added | Percent identity of least-conserved window included | Total number of windows included after adding ties | Coverage of oryLat04 after including ties |
|---|---|---|---|---|---|
| xipCou01 | 100 | 7,460 | 69% | 15,919,720 | 5.00% |
| xipHel01 | 10 | 9,957,730 | 80% | 31,189,076 | 6.43% |
| xipHel01 | 25 | 1,889,845 | 80% | 22,959,004 | 5.30% |
| xipHel01 | 50 | 758,896 | 76% | 20,661,567 | 5.14% |
| xipHel01 | 100 | 99,664 | 64% | 16,301,809 | 5.02% |
| xipMac02 | 10 | 9,027,389 | 90% | 27,849,819 | 6.81% |
| xipMac02 | 25 | 244,169 | 88% | 18,078,229 | 5.05% |
| xipMac02 | 50 | 139,890 | 84% | 17,469,325 | 5.03% |
| xipMac02 | 100 | 802,355 | 78% | 15,399,347 | 5.21% |

**Table S5:** Oligonucleotides used in this study.

| Oligo ID | Oligo Sequence | Description | Species |
|---|---|---|---|
| Faf1CONDEL_ssODN | 5' G*G*CTCGCTTAAACCAGCAGTCAAAGCCCAAC-TACTGCTCTATTTCTGTGGGAAATGTT*C*G 3' | ssODN for pCONDEL.1189 CRISPR; *=phosphorothioate bond | Threespine Stickleback |
| sgRNAscaffoldOligo | 5' GATCCGCACCGACTCGGTGCCACTTTTTCAAGTTGATAACG-GACTAGCCTTATTTTAACTTGCTATTTCTAGCTCTAAAAC 3' | for 2-oligo PCR to generate sgRNA template | n/a |
| 129,589.F_y | CCTTTCACTGTGYAATCAGCTC | amplification, sequencing primer | [multiple species] |
| 129,611.R_r | GAGCTGATTRCACAGTGAAAGG | amplification, sequencing primer | [multiple species] |
| dicLab-1189seq-R2 | CTGATGATCCCCAGCTCCTG | amplification, sequencing primer | European Seabass |
| dicLab-1189seq-R3 | AGGCATCCGTGTTTGAAAGC | amplification, sequencing primer | European Seabass |
| dicLab-CONDEL1189-F1 | TGCCCTGTGCTTTACCCTTT | amplification, sequencing primer | European Seabass |
| dicLab-CONDEL1189-R1 | CTGCAAGGTGTCTCCAGAGG | amplification, sequencing primer | European Seabass |
| dicLab-PelA-F0 | GGTGCCAGGAAGAGTAGCTG | amplification, sequencing primer | European Seabass |
| dicLab-PelA-F1 | CCCTGTGTCATGTCTAAGGCA | amplification, sequencing primer | European Seabass |
| dicLab-PelA-F3 | GTGTCCTATCCATCAGCTGT | amplification, sequencing primer | European Seabass |
| dicLab-PelA-F4 | TACTGTCTGTGATCGCTCTC | amplification, sequencing primer | European Seabass |
| dicLab-PelA-F5 | CCTGTGTGCTGTCAGTGGTA | amplification, sequencing primer | European Seabass |
| dicLab-PelA-pGL4.23-fwd | AGCTCGCTAGCCTCGAGGATTTGAGAGAGGCCTGGAGAAG | primer for Gibson Assembly into pGL4.23 vector | European Seabass |
| dicLab-PelA-pGL4.23-rev | TATATACCCTCTAGTGTCTAGTCCCCTCCTCTATTTTTAA | primer for Gibson Assembly into pGL4.23 vector | European Seabass |
| dicLab-PelA-R1 | GGGGGATCCTGCACTTTGAA | amplification, sequencing primer | European Seabass |
| dicLab-PelA-R2 | ACCCAAGTGTGTGTGTGTGT | amplification, sequencing primer | European Seabass |
| dicLab-PelA-R3 | TGCCTTAGACATGACACAGGG | amplification, sequencing primer | European Seabass |
| dicLab-PelA-R3.5 | CAGCTACTCTTCCTGGCACC | amplification, sequencing primer | European Seabass |
| dicLab-PelA-R4 | ACAATGCACCGTACAGCTCA | amplification, sequencing primer | European Seabass |
| dicLab-PelA-R5 | TCAAACGCATCAGATGGCCT | amplification, sequencing primer | European Seabass |
| dicLab-PelAseq-F2 | ACCTCAGGTCACAGGTTTTCT | amplification, sequencing primer | European Seabass |
| dicLab-PelAseq-R2 | AGTGGCAACGTACAGCAAGA | amplification, sequencing primer | European Seabass |
| FAF1-ex5-R1 | ACTGTGCTGCCTTCCTCCA | amplification, sequencing primer | Threespine Stickleback |
| FAF1-in1-F1 | CCTGTTGTGGGTTGGTACCA | amplification, sequencing primer | Threespine Stickleback |
| FAF1-in2-F1 | CGTGCTTGGCTTTCAGCATT | amplification, sequencing primer | Threespine Stickleback |
| FAF1-in2-R2 | TAGGAACTTGGCGGACACAC | amplification, sequencing primer | Threespine Stickleback |
| FAF1-in3-F2 | TGTCTCCTTCAGTGTTCGGT | amplification, sequencing primer | Threespine Stickleback |
| FAF1-in3-R1 | CAACCACTGTAATCATGTCGCT | amplification, sequencing primer | Threespine Stickleback |
| fr-CONDEL1189-F1 | AAGGTCTCCCCAGGTGGATT | amplification, sequencing primer | Japanese Puffer |
| fr-CONDEL1189-R1 | GGATTCCTGTGCTTCAGCCT | amplification, sequencing primer | Japanese Puffer |

*Continued on next page*

Table S5 – continued from previous page

| Oligo ID | Oligo Sequence | Description | Species |
|---|---|---|---|
| fr-CONDEL1189seq-F2 | TAAGCATTGACCTCTGCTGG | amplification, sequencing primer | Japanese Puffer |
| fr-CONDEL1189seq-R2 | AACGAGCCACTCTACCATGC | amplification, sequencing primer | Japanese Puffer |
| fr3-PelA-F2 | TCACTTCGCAGCTCGTATCC | amplification, sequencing primer | Japanese Puffer |
| fr3-PelA-fwdAnchor | TAATGAAAAGGCCTTAAATCCCTCT | amplification, sequencing primer | Japanese Puffer |
| fr3-PelA-revAnchor | TGTCACCCCAGAGTTATAAAAACTG | amplification, sequencing primer | Japanese Puffer |
| fr3-PelAseq-F1 | GGGTTCGGCCTGGTGATTAA | amplification, sequencing primer | Japanese Puffer |
| fr3-PelAseq-R1 | TGACCCCAAGCTCCATTACG | amplification, sequencing primer | Japanese Puffer |
| gasAcu-CONDEL1189-F1 | GCGTTGGAATAAGCAGCGAG | amplification, sequencing primer | Threespine Stickleback |
| gasAcu-CONDEL1189-R1 | GCAAGGTGTTAGCAGAGGGT | amplification, sequencing primer | Threespine Stickleback |
| gasAcu-CONDEL1189-R2 | ACACAGACTGGTTTGTGGTCCA | amplification, sequencing primer | Threespine Stickleback |
| gasAcu-CONDEL1189seq-F2 | CACAGCAATGAACGGCACAA | amplification, sequencing primer | Threespine Stickleback |
| gasAcu-CONDEL1189seq-R2 | TGTTGGAGGAGCCGATGTTC | amplification, sequencing primer | Threespine Stickleback |
| gasAcu-crispr1189-F1 | CCTACTCCTTGTCTCGGCCAAA | amplification, sequencing primer | Threespine Stickleback |
| gasAcu-crispr1189-R2 | ACACAGACTGGTTTGTGGTCCA | amplification, sequencing primer | Threespine Stickleback |
| HC M13 forward | GTAAAACGACGGCCAGTGAA | vector backbone sequencing primer | n/a [vector backbone] |
| HC M13 reverse | GGAAACAGCTATGACCATGA | vector backbone sequencing primer | n/a [vector backbone] |
| HC Sp6 promoter primer | CTATTTAGGTGACACTATAG | vector backbone sequencing primer | n/a [vector backbone] |
| HC T7 promoter primer | TAATACGACTCACTATAGGG | vector backbone sequencing primer | n/a [vector backbone] |
| HC_pT2HE_rev | CATATAGACAAAACATGCTCGTTC | vector backbone sequencing primer | n/a [vector backbone] |
| hipCom-CONDEL1189-F1 | TTTGAAGAGTGGAGGTGCCC | amplification, sequencing primer | Lined Seahorse |
| hipCom-CONDEL1189-R1 | TGAAGGTGTCCCCAGAAGGT | amplification, sequencing primer | Lined Seahorse |
| hipCom-PelA-F1 | ATCGAGTCAACCAAACCGCT | amplification, sequencing primer | Lined Seahorse |
| hipCom-PelA-R1 | GGCCTGACACTGTCCAGAAG | amplification, sequencing primer | Lined Seahorse |
| hipEre-PelA-F1 | CCCCCTAGTGACCAAGCAAG | amplification, sequencing primer | Lined Seahorse |
| hipEre-PelAseq-F2 | ATCGGCAAACGTTTTGCTTGCTC | amplification, sequencing primer | Lined Seahorse |
| hipEre-PelAseq-F3 | GACACTCGCCTCTTCAATAACC | amplification, sequencing primer | Lined Seahorse |
| hipEre-PelAseq-R1 | CAGTGAGCCAAGTCCATCAA | amplification, sequencing primer | Lined Seahorse |
| larCro-CONDEL1189-F1 | AGGGATGAGGATGAGCAGGA | amplification, sequencing primer | Yellow Croaker |
| larCro-CONDEL1189-F2 | GAGCAGGAGAAATGTTAATGATCAAT | amplification, sequencing primer | Yellow Croaker |
| larCro-CONDEL1189-F3 | CCAGCTCCAGTGCAGAAAGA | amplification, sequencing primer | Yellow Croaker |
| larCro-CONDEL1189-F4 | GCCACTGGATCACTGTCACA | amplification, sequencing primer | Yellow Croaker |
| larCro-CONDEL1189-F5 | TGGAGAGGAGCAATGCACAG | amplification, sequencing primer | Yellow Croaker |
| larCro-CONDEL1189-F6 | GTCATTCACAGTCCGAGGCA | amplification, sequencing primer | Yellow Croaker |
| larCro-CONDEL1189-F7 | AAAAGGTTCAGTGCCAGGCA | amplification, sequencing primer | Yellow Croaker |
| larCro-CONDEL1189-F8 | GCAACAGATTGGACCCCTGT | amplification, sequencing primer | Yellow Croaker |
| larCro-CONDEL1189-F9 | TGCCATGAAAATGGAGCCCT | amplification, sequencing primer | Yellow Croaker |
| larCro-CONDEL1189-R1 | GCCTTTGTCAGAGAAGCGCT | amplification, sequencing primer | Yellow Croaker |
| larCro-CONDEL1189-R2 | CAACACAGCAGGTATGGCAC | amplification, sequencing primer | Yellow Croaker |
| larCro-CONDEL1189-R3 | ACAGGGGTCCAATCTGTTGC | amplification, sequencing primer | Yellow Croaker |
| larCro-CONDEL1189-R4 | GCATCAGCCTTTTCACAGCC | amplification, sequencing primer | Yellow Croaker |
| larCro-CONDEL1189-R5 | CCCATCTGCCAATGGTCACT | amplification, sequencing primer | Yellow Croaker |
| larCro-CONDEL1189-R6 | TGTGACAGTGATCCAGTGGC | amplification, sequencing primer | Yellow Croaker |
| larCro-PelA-3prime | CCCTACTCCTCTATTTTTAA | amplification, sequencing primer | Yellow Croaker |
| larCro-PelA-5prime | TTGAGAGAGGCCTGGAGAGG | amplification, sequencing primer | Yellow Croaker |
| larCro-PelA-F3.1 | GATTTACATTTGCCAGCTGG | amplification, sequencing primer | Yellow Croaker |
| larCro-PelA-F3.2 | TAAAAGACTACACGGTCTCT | amplification, sequencing primer | Yellow Croaker |
| larCro-PelA-F4.1 | CTCTCTCCTTCCAAGTATTCCAAG | amplification, sequencing primer | Yellow Croaker |
| larCro-PelA-innerFwdAnchor | ATTTGACCTTTGTCCCCTCCC | amplification, sequencing primer | Yellow Croaker |
| larCro-PelA-innerRevAnchor | AAAAGTGATGTAAACATTGAGAGAGGC | amplification, sequencing primer | Yellow Croaker |
| larCro-PelA-outerFwdAnchor | TAATGATAAGGCCTTAAATCCCTCTG | amplification, sequencing primer | Yellow Croaker |
| larCro-PelA-outerRevAnchor | GTGTGCAGTACTTGGGCTCT | amplification, sequencing primer | Yellow Croaker |
| larCro-PelA-pGL4.23-fwd | AGCTCGCTAGCCTCGAGGATTTGAGAGAGGCCTGGAGAGG | primer for Gibson Assembly into pGL4.23 vector | Yellow Croaker |
| larCro-PelA-pGL4.23-rev | TATATACCCTCTAGTGTCTAGTCCCCTCCCCCCTACTCCT | primer for Gibson Assembly into pGL4.23 vector | Yellow Croaker |
| larCro-PelA-pT2HE-GA-fwd | GGCCCGATATCACCGGGGGATTGAGAGAGGCCTGGAGAGG | primer for Gibson Assembly into pT2HE vector | Yellow Croaker |

*Continued on next page*

**Table S5 – continued from previous page**

| Oligo ID | Oligo Sequence | Description | Species |
|---|---|---|---|
| larCro-PelA-pT2HE-GA-rev | CAAGCGACACCCCTGAAGGAGTCCCCTCCCCCCTACTCCT | primer for Gibson Assembly into pT2HE vector | Yellow Croaker |
| larCro-PelA-R1 | TGACAAGCCCACACACTGTT | amplification, sequencing primer | Yellow Croaker |
| larCro-PelA-R1.5 | GTCTGGGCTCTGCTCGTTTG | amplification, sequencing primer | Yellow Croaker |
| larCro-PelA-R1.75 | CACAATTAATCAGTGGGCCAAA | amplification, sequencing primer | Yellow Croaker |
| larCro-PelA-R2 | GGCCAAATGCACAATGGGTT | amplification, sequencing primer | Yellow Croaker |
| larCro-PelA-R3 | CGCAAAGCCCTCCTACTGAA | amplification, sequencing primer | Yellow Croaker |
| larCro-PelA-R4 | TGAGGCAAGGGGGTTGTAAC | amplification, sequencing primer | Yellow Croaker |
| larCro-PelA-R4.1 | CTTGGAATACTTGGAAGGAGAGAG | amplification, sequencing primer | Yellow Croaker |
| larCro-PelA-R5 | ATTCTCTTTGACCCTGCCGG | amplification, sequencing primer | Yellow Croaker |
| larCro-PelA-R6 | TCATTTAGAAATCTTGGATATCTAA | amplification, sequencing primer | Yellow Croaker |
| larCro-PelAseq-F0 | CACAGCCACGTTCCAAACAC | amplification, sequencing primer | Yellow Croaker |
| larCro-PelAseq-F0.5 | TTTGGCCCACTGATTAATTGTG | amplification, sequencing primer | Yellow Croaker |
| larCro-PelAseq-F1 | AACCCATTGTGCATTTGGCC | amplification, sequencing primer | Yellow Croaker |
| larCro-PelAseq-F2 | TTCAGTAGGAGGGCTTTGCG | amplification, sequencing primer | Yellow Croaker |
| larCro-PelAseq-F2.5 | GAGGTGTCATTTTCTGTGAGGAG | amplification, sequencing primer | Yellow Croaker |
| larCro-PelAseq-F3 | AACAGTGTGTGGGCTTGTCA | amplification, sequencing primer | Yellow Croaker |
| larCro-PelAseq-F4 | GTTACAACCCCCTTGCCTCA | amplification, sequencing primer | Yellow Croaker |
| larCro-PelAseq-F5 | CCGGCAGGGTCAAAGAGAAT | amplification, sequencing primer | Yellow Croaker |
| larCro-PelAseq-F6 | GTGAAAGGATCCCTCGGTGT | amplification, sequencing primer | Yellow Croaker |
| larCro-PelAseq-F7 | GCGGTGAGGTGAATACGGAT | amplification, sequencing primer | Yellow Croaker |
| larCro-PelAseq-F8 | CCACCTTCCCAGATCTCAGC | amplification, sequencing primer | Yellow Croaker |
| larCro0-PelA-R3.1 | CTCCTCACAGAAAATGACACCTC | amplification, sequencing primer | Yellow Croaker |
| luc2-forMinP | GTCCACCTCGATATGTGCGT | amplification, sequencing primer | Yellow Croaker |
| luc2rev | CCCTTCTTAATGTTTTTGGC | vector backbone sequencing primer | n/a [vector backbone] |
| minP-rev | ATTGCCAAGCTGGAAGTCGA | vector backbone sequencing primer | n/a [vector backbone] |
| molMol-CONDEL1189-F1 | GGTACAGATGGCGTTCGGAA | amplification, sequencing primer | Ocean Sunfish |
| molMol-CONDEL1189-R1 | AGGGGTAAGGATCAGCAGGA | amplification, sequencing primer | Ocean Sunfish |
| molMol-PelA-F1 | GCTGCAGCGTTCTTTGATGT | amplification, sequencing primer | Ocean Sunfish |
| molMol-PelA-F6 | TTTTCCACACACGCCTAGCT | amplification, sequencing primer | Ocean Sunfish |
| molMol-PelA-F7 | TCCAATCAAATTCCATTCAGCCG | amplification, sequencing primer | Ocean Sunfish |
| molMol-PelA-fwdAnchor | TGTCATCCCAGAGTTATAAAAAGTGA | amplification, sequencing primer | Ocean Sunfish |
| molMol-PelA-pGL4.23-fwd | AGCTCGCTAGCCTCGAGGATCTGAGCGGGGCCTGGATAGG | primer for Gibson Assembly into pGL4.23 vector | Ocean Sunfish |
| molMol-PelA-pGL4.23-rev | TATATACCCTCTAGTGTCTAGTCCCGTCTATTTTTAAAAC | primer for Gibson Assembly into pGL4.23 vector | Ocean Sunfish |
| molMol-PelA-R1 | CCTGTGTCATTGGGTCCACA | amplification, sequencing primer | Ocean Sunfish |
| molMol-PelA-R5 | AAGTCAGTGATCCAGCTGGC | amplification, sequencing primer | Ocean Sunfish |
| molMol-PelA-R6 | GGGGTGGCATGTTTGTTGAG | amplification, sequencing primer | Ocean Sunfish |
| molMol-PelA-revAnchor | TAATGGTAAGGCCTTAAATCCCCC | amplification, sequencing primer | Ocean Sunfish |
| molMol-PelAseq-F2 | CTCAACAAACATGCCACCCC | amplification, sequencing primer | Ocean Sunfish |
| molMol-PelAseq-F3 | GTGCTTCTCCTGCAAAAGGC | amplification, sequencing primer | Ocean Sunfish |
| molMol-PelAseq-F4 | GCCAGCTGGATCACTGACTT | amplification, sequencing primer | Ocean Sunfish |
| molMol-PelAseq-F5 | TCTGGGCTTTTTGGGTGTGT | amplification, sequencing primer | Ocean Sunfish |
| molMol-PelAseq-R2 | TCCACTCATGTGATGCCCTG | amplification, sequencing primer | Ocean Sunfish |
| molMol-PelAseq-R3 | CACTGATTGTAACAGTTGTCGC | amplification, sequencing primer | Ocean Sunfish |
| molMol-PelAseq-R4 | GCAGCACTTTGGAGGTCCTC | amplification, sequencing primer | Ocean Sunfish |
| monAlb-1189-F2 | CTGCTCATATCCAGGCCCAG | amplification, sequencing primer | Rice Eel |
| monAlb-1189-R2 | CCGAGTCAAAACGGTCTCCA | amplification, sequencing primer | Rice Eel |
| monAlb-1189seq-F1 | GAGACGAGGGTTCCATGCAA | amplification, sequencing primer | Rice Eel |
| monAlb-1189seq-R1 | ACCCTCTTCTCTGCCTTCCT | amplification, sequencing primer | Rice Eel |
| monAlb-CONDEL1189-F1 | GCAGCAAGAGGAGAGATGCA | amplification, sequencing primer | Rice Eel |
| monAlb-CONDEL1189-R1 | GGGCAAGGATCAGCAAGAGA | amplification, sequencing primer | Rice Eel |
| oryLat-CONDEL1189-F1 | ATACTGACAGTGATGCGGCA | PCR amplification, sequencing primer | Japanese Medaka |
| oryLat-CONDEL1189-F2 | ACCGCAGTGAAATCAGCCTT | PCR amplification, sequencing primer | Japanese Medaka |
| oryLat-CONDEL1189-F3 | CAATGCAACGGAGACACACC | PCR amplification, sequencing primer | Japanese Medaka |
| oryLat-CONDEL1189-F4 | TTTCCTCCCTCTCGCCTTTCTG | PCR amplification, sequencing primer | Japanese Medaka |

*Continued on next page*

**Table S5 – continued from previous page**

| Oligo ID | Oligo Sequence | Description | Species |
|---|---|---|---|
| oryLat-CONDEL1189-R1 | GGATCAGGGTGGCTTCAAGTGA | PCR amplification, sequencing primer | Japanese Medaka |
| oryLat-CONDEL1189-R2 | CCGTGCCTCTCGTGGTAAAT | PCR amplification, sequencing primer | Japanese Medaka |
| oryLat-CONDEL1189-R3 | TTTAGTGCTCTCCGTGCCTC | PCR amplification, sequencing primer | Japanese Medaka |
| oryLat-CONDEL1189-R4 | TCATGGATCAGGGTGGCTTCAA | PCR amplification, sequencing primer | Japanese Medaka |
| oryLat-CONDEL1189-preNheIfwd3 | AGATAGGCCCCTTACGTACGCTCAATGCAACGGAGACACACC | primer for Gibson Assembly into pT2HE vector | Japanese Medaka |
| oryLat-CONDEL1189-postNheIrev1 | CCGGTGATATCGGGCCCGCTGGATCAGGGTGGCTTCAAGTGA | primer for Gibson Assembly into pT2HE vector | Japanese Medaka |
| oryLat-DEL963-F1 | CACATTTCTGCAAGGTGCCC | PCR amplification, sequencing primer | Japanese Medaka |
| oryLat-DEL963-F3 | TGGACACTTGTACAGGCCACAT | PCR amplification, sequencing primer | Japanese Medaka |
| oryLat-DEL963-F4 | TGACAGATTCCCATTTGCATGTGT | PCR amplification, sequencing primer | Japanese Medaka |
| oryLat-DEL963-F5 | AGTCGTCATGAGGATTCCCTTT | PCR amplification, sequencing primer | Japanese Medaka |
| oryLat-DEL963-F6 | AGGGGACTGAAAGAGGAAAGTTGT | PCR amplification, sequencing primer | Japanese Medaka |
| oryLat-DEL963-F7 | CCGTGTTCATTTCAGGGGCAAT | PCR amplification, sequencing primer | Japanese Medaka |
| oryLat-DEL963-F8 | TCTGGACCGTTCGAATCAAGCT | PCR amplification, sequencing primer | Japanese Medaka |
| oryLat-DEL963-F9 | TGTGGAGGTTTGACTGACTGCT | PCR amplification, sequencing primer | Japanese Medaka |
| oryLat-DEL963-R10 | ATTGCCCCTGAAATGAACACGG | PCR amplification, sequencing primer | Japanese Medaka |
| oryLat-DEL963-R11 | ACAACTTTCCTCTTTCAGTCCCCT | PCR amplification, sequencing primer | Japanese Medaka |
| oryLat-DEL963-R3 | CTTTAGCCTCTGTCAGAACGCG | PCR amplification, sequencing primer | Japanese Medaka |
| oryLat-DEL963-R4 | CCCTCCATGTGGCCTGTACAAG | PCR amplification, sequencing primer | Japanese Medaka |
| oryLat-DEL963-R5 | TGAGCTGGAAATGGAGACTCGT | PCR amplification, sequencing primer | Japanese Medaka |
| oryLat-DEL963-R8 | AGCTTGATTCGAACGGTCCAGA | PCR amplification, sequencing primer | Japanese Medaka |
| oryLat-DEL963-R9 | GGGTCCTTCCAACTGTTTGCAC | PCR amplification, sequencing primer | Japanese Medaka |
| oryLat-PelA-F1 | GGGGAATCCTGCACTTTGAA | amplification, sequencing primer | Japanese Medaka |
| oryLat-PelA-F5 | CTCTAGTCTGATATTGAAGCGTATT | amplification, sequencing primer | Japanese Medaka |
| oryLat-PelA-F6 | TTAAGCTCCTGATCGCCGCT | amplification, sequencing primer | Japanese Medaka |
| oryLat-PelA-fwdAnchor | TGTCACTCCAGAGTTCTAAAGAGT | amplification, sequencing primer | Japanese Medaka |
| oryLat-PelA-pGL4.23-fwd | AGCTCGCTAGCCTCGAGGATCTGAAAGTGAGGCCAGGAGA | primer for Gibson Assembly into pGL4.23 vector | Japanese Medaka |
| oryLat-PelA-pGL4.23-rev | TATATACCCTCTAGTGTCTAGTCCTCCTTTTTTCTCTATT | primer for Gibson Assembly into pGL4.23 vector | Japanese Medaka |
| oryLat-PelA-R1 | CGTTCTTCCTTCATCGCTGC | amplification, sequencing primer | Japanese Medaka |
| oryLat-PelA-R4 | GGTCTGCAGGTGACACGTAA | amplification, sequencing primer | Japanese Medaka |
| oryLat-PelA-revAnchor | GATAAAGCCCTAAATCCTTCTGTAAA | amplification, sequencing primer | Japanese Medaka |
| oryLat-PelAseq-F2 | TGCGTCTGTTCGTGCAAAAG | amplification, sequencing primer | Japanese Medaka |
| oryLat-PelAseq-F3 | CATGCTTACAGCCCCTCACA | amplification, sequencing primer | Japanese Medaka |
| oryLat-PelAseq-F4 | TTACGTGTCACCTGCAGACC | amplification, sequencing primer | Japanese Medaka |
| oryLat-PelAseq-R2 | ACCCTCGGTTTAAGCATTAAGGT | amplification, sequencing primer | Japanese Medaka |
| oryLat-PelAseq-R3 | AGCCCATCGGAACCTTTCAG | amplification, sequencing primer | Japanese Medaka |
| PelA 129,078.F | CATCACCGAGCCGCTTTGAT | amplification, sequencing primer | Threespine Stickleback |
| PelA 129,096.F | ATGTGGGCCTAATATGGCTTG | amplification, sequencing primer | Threespine Stickleback |
| PelA 129,165.R | TCTGGACAGCGTCAGGCAAC | amplification, sequencing primer | Threespine Stickleback |
| PelA 129,231.F | GTCTGCAGCTGTCATCCTG | amplification, sequencing primer | Threespine Stickleback |
| PelA 129,237.F | CTGTCATCCTGGAGTTATAAAAC | amplification, sequencing primer | Threespine Stickleback |
| PelA 129,262.F | CATCCTGGAGTTATAAAACGTG | amplification, sequencing primer | Threespine Stickleback |
| PelA 129,282.F | GATGTAAACATTGAGAGGGTC | amplification, sequencing primer | Threespine Stickleback |
| PelA 129,334.F | CTGTCATGGAGCACCACCTC | amplification, sequencing primer | Threespine Stickleback |
| PelA 129,345.F | CACCACCTCGTTCCTGAAGG | amplification, sequencing primer | Threespine Stickleback |
| PelA 129,345.R | CCTTCAGGAACGAGGTGGTG | amplification, sequencing primer | Threespine Stickleback |
| PelA 129,410.F | CCATTTACTAAAAATGCTCAACTC | amplification, sequencing primer | Threespine Stickleback |
| PelA 129,433.R | GAGTTGAGCATTTTTAGTAAATGG | amplification, sequencing primer | Threespine Stickleback |
| PelA 129,451.F | CTCGATTTCCATCACGTTGTT | amplification, sequencing primer | Threespine Stickleback |
| PelA 129,490.F | GCACATGAAGGATCATTAACAG | amplification, sequencing primer | Threespine Stickleback |
| PelA 129,504.R | GTGCACCTGAATTACTGTTAATG | amplification, sequencing primer | Threespine Stickleback |
| PelA 129,508.F | CAGTAATTCAGGTGCACAAAC | amplification, sequencing primer | Threespine Stickleback |
| PelA 129,560.F | CAGCCTGATGTGCAGCACAC | amplification, sequencing primer | Threespine Stickleback |
| PelA 129,656.R | AGCTTATCTCGGCTGTTTATGT | amplification, sequencing primer | Threespine Stickleback |
| PelA 129,689.F | GTCGAAGCAAAGAGGCGAG | amplification, sequencing primer | Threespine Stickleback |

*Continued on next page*

**Table S5 – continued from previous page**

| Oligo ID | Oligo Sequence | Description | Species |
|---|---|---|---|
| PelA 129,750.R | GATATCTGAATGTTTATGAATAACA | amplification, sequencing primer | Threespine Stickleback |
| PelA 129,754.F | CCAACAAGAAACTGTTCAAATG | amplification, sequencing primer | Threespine Stickleback |
| PelA 129,918.F | TCAGGGCCGGACCATCTAA | amplification, sequencing primer | Threespine Stickleback |
| PelA 129,968.R | GCAGAGTTCTAAAGTGGTCG | amplification, sequencing primer | Threespine Stickleback |
| PelA 129,976.R | GACCTTGTGCAGAGTTCTAAAG | amplification, sequencing primer | Threespine Stickleback |
| PelA 129,985.R | ATGCATTAGGACCTTGTGCAG | amplification, sequencing primer | Threespine Stickleback |
| PelA 129,987.F | CTGTTTGACCTCGCCGGAG | amplification, sequencing primer | Threespine Stickleback |
| PelA 129,993.R | CAAACAGAATGCATTAGGACC | amplification, sequencing primer | Threespine Stickleback |
| PelA 129,999.R | CGAGGTCAAACAGAATGCATTAG | amplification, sequencing primer | Threespine Stickleback |
| PelA 129.937.F | CTCTTCCACTGATTGTTATG | amplification, sequencing primer | Threespine Stickleback |
| PelA 130,000.F | CCGGAGTAAATCAAATACTGG | amplification, sequencing primer | Threespine Stickleback |
| PelA 130,020.R | CCAGTATTTGATTTACTCCGG | amplification, sequencing primer | Threespine Stickleback |
| PelA 130,164.F | CACGCTAGACACAAGGAAGG | amplification, sequencing primer | Threespine Stickleback |
| PelA 130,250.R | GTGACCACAACAATCCGTGG | amplification, sequencing primer | Threespine Stickleback |
| PelA 130,269.F | CCACGGATTGTTGTGGTCAC | amplification, sequencing primer | Threespine Stickleback |
| PelA 130,322.F | AGCTAGCCGCTAACAGGTAG | amplification, sequencing primer | Threespine Stickleback |
| PelA 130,377.F | AGTAGCGTTCAACTCTTTCTAG | amplification, sequencing primer | Threespine Stickleback |
| PelA 130,398.R | CTAGAAAGAGTTGAACGCTACT | amplification, sequencing primer | Threespine Stickleback |
| PelA 130,485.R | GGTGGTTATTTAAATAGAGACAATA | amplification, sequencing primer | Threespine Stickleback |
| PelA 130,520.R | GACATGCTGGTCTATCAGAC | amplification, sequencing primer | Threespine Stickleback |
| PelA 130,547.F | GTGTTCTTCATAATACAGAATCAGCATC | amplification, sequencing primer | Threespine Stickleback |
| PelA 130,577.R | GAAGATGCTGATTCTGTATTATG | amplification, sequencing primer | Threespine Stickleback |
| PelA 130,625.R | CACGGAGGACGTCTTTCAGG | amplification, sequencing primer | Threespine Stickleback |
| PelA 130,634.R | GACAACTCGCACGGAGGAC | amplification, sequencing primer | Threespine Stickleback |
| PelA 130,643.F | GTACTGCGATAGATCTGAGG | amplification, sequencing primer | Threespine Stickleback |
| PelA 130,662.R | CCTCAGATCTATCGCAGTAC | amplification, sequencing primer | Threespine Stickleback |
| PelA 130,665.R | GGACCTCAGATCTATCGCAG | amplification, sequencing primer | Threespine Stickleback |
| PelA 130,757.R | CTCCTGCTGGAGGACCTTAG | amplification, sequencing primer | Threespine Stickleback |
| PelA 130,778.R | GTTTGTCAGCAGAGAGGTTTC | amplification, sequencing primer | Threespine Stickleback |
| PelA 130,877.F | CACGCCCCGTCCTCCCGA | amplification, sequencing primer | Threespine Stickleback |
| PelA 130,920.R | TTTTTATTTTGATATGTCTCCTGC | amplification, sequencing primer | Threespine Stickleback |
| PelA 130,943.R | AAATCCCTCTGTAGATTTGACC | amplification, sequencing primer | Threespine Stickleback |
| PelA 130,957.R | CGATAAGGCCTTAAATCCCTC | amplification, sequencing primer | Threespine Stickleback |
| PelA 131,454.R | GAAATGAGAACATTTCACATCTAC | amplification, sequencing primer | Threespine Stickleback |
| PelA 131,480.F | GTCTGAAATATGTTATAGGTGC | amplification, sequencing primer | Threespine Stickleback |
| PelA 131,644.R | TCATGTTGTAGAATGAATTGAGC | amplification, sequencing primer | Threespine Stickleback |
| PelA 131,675.F | CCTGCAGTAAATAACTAGGAG | amplification, sequencing primer | Threespine Stickleback |
| PelA 131,713.R | ATAGTTTAGTATAGTATACTCCTAG | amplification, sequencing primer | Threespine Stickleback |
| PelA 131,758.R | TCTCTCAGCGGAGAAATCCG | amplification, sequencing primer | Threespine Stickleback |
| PelA 132,268.R | AGCTTCGTCACGCCACCTG | amplification, sequencing primer | Threespine Stickleback |
| PelA_insertion_fwd1 | GCGTGTGATTGCCAGAGACATT | amplification, sequencing primer | Threespine Stickleback |
| PelA_insertion_fwd2 | GTGTGGCTGGTCTCGTATCATA | amplification, sequencing primer | Threespine Stickleback |
| PelA_insertion_rev1 | GAATGTTAATAATTTAAGGCTGT | amplification, sequencing primer | Threespine Stickleback |
| PelA_insertion_rev1fwd | ACAGCCTTAAATTATTAACATTC | amplification, sequencing primer | Threespine Stickleback |
| PelA_insertion_rev2 | ATAATTTAAGGCTGTCGGGGGT | amplification, sequencing primer | Threespine Stickleback |
| PelA_insertion_rev3 | AATGTCTCTGGCAATCACACGC | amplification, sequencing primer | Threespine Stickleback |
| PelA_insertion_rev4 | CTCATGGGGGAATGTTAATA | amplification, sequencing primer | Threespine Stickleback |
| percomorph-COI-fwd | ttctccaccaaccacaargayatygg | amplification, sequencing primer | [multiple species] |
| percomorph-COI-rev | cacctcagggtgtccgaaraaycaraa | amplification, sequencing primer | [multiple species] |
| RVprimer3 | TAGCAAAATAGGCTGTCCC | vector backbone sequencing primer | n/a [vector backbone] |
| SALR-PelA-pGL4.23-fwd | AGCTCGCTAGCCTCGAGGATTTGAGAGGGTCTGGAGGAGC | primer for Gibson Assembly into pGL4.23 vector | Threespine Stickleback |
| SALR-PelA-pGL4.23-rev | TATATACCCTCTAGTGTCTATTTATTTTGATATGTCTCCT | primer for Gibson Assembly into pGL4.23 vector | Threespine Stickleback |
| SALR-PelA-pT2HE-GA-fwd | GGCCCGATATCACCGGGGGATTGAGAGGGTCTGGAGGAGC | primer for Gibson Assembly into pT2HE vector | Threespine Stickleback |
| SALR-PelA-pT2HE-GA-rev | CAAGCGACACCCCTGAAGGATTTATTTTGATATGTCTCCT | primer for Gibson Assembly into pT2HE vector | Threespine Stickleback |

*Continued on next page*

Table S5 – continued from previous page

| Oligo ID | Oligo Sequence | Description | Species |
|---|---|---|---|
| seqLuc2fwd | TAGACACTAGAGGGTATATAATGGA | vector backbone sequencing primer | n/a [vector backbone] |
| seqLuc2rev | CTGCATTCTAGTTGTGGTTTGTC | vector backbone sequencing primer | n/a [vector backbone] |
| seqLuc2revMid | CCAGTGTCTTACCGGTGTCC | vector backbone sequencing primer | n/a [vector backbone] |
| seqSV40polyA_fwd | GCAAGATCGCCGTGTAATAA | vector backbone sequencing primer | n/a [vector backbone] |
| serDum-CONDEL1189-F1 | CTGCAAGGTGTCCTCAGAGG | amplification, sequencing primer | Greater Amberjack |
| serDum-CONDEL1189-F2 | CCCACCTGCCTCAAAACTGA | amplification, sequencing primer | Greater Amberjack |
| serDum-CONDEL1189-F3 | GCACGCACAGGTTGAATTGT | amplification, sequencing primer | Greater Amberjack |
| serDum-CONDEL1189-F4 | TTCCTCCTGAACTGGGGACT | amplification, sequencing primer | Greater Amberjack |
| serDum-CONDEL1189-R1 | TTTGAAGAGTGGAGGTGCCC | amplification, sequencing primer | Greater Amberjack |
| serDum-CONDEL1189-R2 | CCCTGTGCTTTAGCCTCTGT | amplification, sequencing primer | Greater Amberjack |
| serDum-CONDEL1189-R3 | AGCTCCCGTGTTGATCACTG | amplification, sequencing primer | Greater Amberjack |
| serDum-PelA-F0 | AATCTTTACCAGGGGCCTGC | amplification, sequencing primer | Greater Amberjack |
| serDum-PelA-F0.5 | TGCCAAGTTCCCCACATCAA | amplification, sequencing primer | Greater Amberjack |
| serDum-PelA-F1 | TCCTTTTTCTGGGCTCACGG | amplification, sequencing primer | Greater Amberjack |
| serDum-PelA-pGL4.23-fwd | AGCTCGCTAGCCTCGAGGATTTGGGAGAGGCCTGGAGAAA | primer for Gibson Assembly into pGL4.23 vector | Greater Amberjack |
| serDum-PelA-pGL4.23-rev | TATATACCCTCTAGTGTCTATTTGTCCCCTCCTCTATTTT | primer for Gibson Assembly into pGL4.23 vector | Greater Amberjack |
| serDum-PelA-R1 | GGGCGATCCTGCTCTTTGAA | amplification, sequencing primer | Greater Amberjack |
| serDum-PelA-R3 | CCGTGAGCCCAGAAAAAGGA | amplification, sequencing primer | Greater Amberjack |
| serDum-PelAseq-F2 | CTGTAGCAGGGTTAGTGCCA | amplification, sequencing primer | Greater Amberjack |
| serDum-PelAseq-R2 | TGGCCAGTAATGGTTGCTGT | amplification, sequencing primer | Greater Amberjack |
| sgRNA-01 | AATTAATACGACTCACTATAGGAATCAATAGACTGGGCCGGTTTTAGAGCTAGAAATAGC | for 2-oligo PCR to generate sgRNA template | Threespine Stickleback |
| sgRNA-02 | AATTAATACGACTCACTATAGGTGCAGCAACTTCCCCAGTGGTTTTAGAGCTAGAAATAG | for 2-oligo PCR to generate sgRNA template | Threespine Stickleback |
| sgRNA-04 | AATTAATACGACTCACTATAGGCCGGATGCTCAACTTCCGGGGTTTTAGAGCTAGAAATA | for 2-oligo PCR to generate sgRNA template | Threespine Stickleback |
| sgRNA-16 | AATTAATACGACTCACTATAGGAAATAGAGCAGTAGAGACAGTTTTAGAGCTAGAAATAG | for 2-oligo PCR to generate sgRNA template | Threespine Stickleback |
| sgRNA-17 | AATTAATACGACTCACTATAGGCAGTCAAAGCCCAAGATCAGTTTTAGAGCTAGAAATAG | for 2-oligo PCR to generate sgRNA template | Threespine Stickleback |
| sgRNA-32 | AATTAATACGACTCACTATAGGTGGTAAGTTCTTTATCTGTGTTTTAGAGCTAGAAATAG | for 2-oligo PCR to generate sgRNA template | Threespine Stickleback |
| sgRNA-35 | AATTAATACGACTCACTATAGGTCATTACAGGCAGCCATCAAGTTTTAGAGCTAGAAATA | for 2-oligo PCR to generate sgRNA template | Threespine Stickleback |
| sgRNA-SLC24A5 | AATTAATACGACTCACTATAGAAGCCGTCGGGGAACTCGGGTTTTAGAGCTAGAAATAGC | for 2-oligo PCR to generate sgRNA template | Threespine Stickleback |
| synSco-CONDEL1189-F1 | AGCACACACCAGGGATGTAC | amplification, sequencing primer | Gulf Pipefish |
| synSco-CONDEL1189-R1 | AGCGCAGACTTTGAAGAGCT | amplification, sequencing primer | Gulf Pipefish |
| synSco-PelA-F1 | TGTTGTTACTGCCCCCTAGC | amplification, sequencing primer | Gulf Pipefish |
| synSco-PelA-fwdAnchor | TGAAGTTAAGGCCCTTAAATCCC | amplification, sequencing primer | Gulf Pipefish |
| synSco-PelA-pGL4.23-fwd | AGCTCGCTAGCCTCGAGGATCAAAAGAAAGAGTCTGGATA | primer for Gibson Assembly into pGL4.23 vector | Gulf Pipefish |
| synSco-PelA-pGL4.23-rev | TATATACCCTCTAGTGTCTAGAACCTTCCCCTCATCCCCT | primer for Gibson Assembly into pGL4.23 vector | Gulf Pipefish |
| synSco-PelA-R1 | GCACTTCATCGCAGTTCTCG | amplification, sequencing primer | Gulf Pipefish |
| synSco-PelA-R1 | GCACTTCATCGCAGTTCTCG | amplification, sequencing primer | Gulf Pipefish |
| synSco-PelA-revAnchor | TGTCATTCCGGAGTTATAAACGT | amplification, sequencing primer | Gulf Pipefish |
| synSco-PelA-seq-R1 | AAGGCGGGATATGCGTCAAA | amplification, sequencing primer | Gulf Pipefish |
| tetNig-CONDEL1189-F1 | TCTCCCCAGGTGCATTAGGA | amplification, sequencing primer | Green Spotted Puffer |
| tetNig-CONDEL1189-R1 | GGATTCCTGTGCTTCAGCCT | amplification, sequencing primer | Green Spotted Puffer |
| tetNig-PelA-3prime | GTCCCGTCTATTTTTAAACC | amplification, sequencing primer | Green Spotted Puffer |
| tetNig-PelA-5prime | CTGAGAAAAGTCTGGAGAAG | amplification, sequencing primer | Green Spotted Puffer |
| tetNig-PelA-fwdAnchor | TGTGACCCCAGAGTTATAAAAACTG | amplification, sequencing primer | Green Spotted Puffer |
| tetNig-PelA-pGL4.23-fwd | AGCTCGCTAGCCTCGAGGATCTGAGAAAAGTCTGGAGAAG | primer for Gibson Assembly into pGL4.23 vector | Green Spotted Puffer |
| tetNig-PelA-pGL4.23-rev | TATATACCCTCTAGTGTCTAGTCCCGTCTATTTTTAAACC | primer for Gibson Assembly into pGL4.23 vector | Green Spotted Puffer |
| tetNig-PelA-pT2HE-GA-fwd | GGCCCGATATCACCGGGGGACTGAGAAAAGTCTGGAGAAG | primer for Gibson Assembly into pT2HE vector | Green Spotted Puffer |
| tetNig-PelA-pT2HE-GA-rev | CAAGCGACACCCCTGAAGGAGTCCCGTCTATTTTTAAACC | primer for Gibson Assembly into pT2HE vector | Green Spotted Puffer |
| tetNig-PelA-R1 | GAGTTCACCGTACAGCCTTC | amplification, sequencing primer | Green Spotted Puffer |
| tetNig-PelA-revAnchor | TGGAAAGGCCTTAAATCCCTCT | amplification, sequencing primer | Green Spotted Puffer |
| tetNig-PelAseq-F1 | TTTGCTGGCTACGTCTCAGG | amplification, sequencing primer | Green Spotted Puffer |

**Table S6:** Sources of images and illustrations in Figure 1.

| Species | Type of Depiction | Artist/Creator | Source of Digital Copy or Tissue | Copyright Holder | Usage Rights |
|---|---|---|---|---|---|
| European Seabass | illustration | G. Cuvier<br>M. Valenciennes | [25] | Public Domain | [not under license] |
| Green Spotted Puffer | photo of stained caudal skeleton (image inset paired with Japanese Puffer) | H. I. Chen<br>D. M. Kingsley | purchased specimen [24] | [authors of this study] | n/a |
| Gulf Pipefish | illustration | J. Tomelleri | [26] | J. Tomelleri | usage rights purchased |
| Japanese Medaka | illustration | D. Jordan, C. Metz | [27] | Public Domain | [not under license] |
| Japanese Puffer | illustration | B. Yau | [28] | B. Yau | [29] |
| Lined Seahorse | radiograph | S. Raredon | [30] | Public Domain | [not under license] |
| Ocean Sunfish | illustration | H. Smith | [31] | Public Domain | [not under license] |
| Ocean Sunfish | illustration (of skeleton) | J. Steenstrup<br>C. Lütken | [15] | Public Domain | [not under license] |
| Rice Eel | illustration | F. Day | [32] | Public Domain | [not under license] |
| Rice Eel | photo of stained caudal skeleton | H. I. Chen<br>D. M. Kingsley | Leo Nico Specimen LGN 12-10 | [authors of this study] | n/a |
| Threespine Stickleback | illustration | E. Edmonson | [33] | NY Department of Environmental Conservation | CC BY-NC-ND 2.0 |
| Tongue Sole | illustration | D. Jordan, C. Metz | [27] | Public Domain | [not under license] |
| Tongue Sole | caudal radiograph | D. Loffler | [34] | Public Domain | [not under license] |
| Yellow Croaker | illustration | B. Yau | [35] | B. Yau | [29] |

[10] T. H. Huxley. Observations on the development of some parts of the skeleton of fishes. *Transactions of The Microscopical Society & Journal*, 7(1):33–46, 1859. ISSN 1365-2818. doi: 10.1111/j.1365-2818.1859.tb04575.x. URL http://onlinelibrary.wiley.com/doi/abs/10.1111/j.1365-2818.1859.tb04575.x.

[11] J. S. Nelson, T. Grande, and M. V. H. Wilson. *Fishes of the world*. John Wiley & Sons, Hoboken, New Jersey, 5th edition, 2016. ISBN 978-1-118-34233-6.

[12] T. Franz-Odendaal and D. Adriaens. Comparative developmental osteology of the seahorse skeleton reveals heterochrony amongst *Hippocampus* sp. and progressive caudal fin loss. *EvoDevo*, 5(45):1–11, 2014. doi: 10.1186/2041-9139-5-45.

[13] Q.-R. Wang, Y.-Y. Ni, L.-M. Lin, and Z.-Y. Wang. Development of the vertebral column and the pectoral and caudal fins in larvae of yellow croaker *Larimichthys crocea* (Richardson). *Acta Hydrobiologica Sinica*, 34(3):467–472, 2010. doi: 10.3724/SP.J.1035.2010.00467.

[14] J. C. Tyler. *Osteology, phylogeny, and higher classification of the fishes of the order Plectognathi (Tetraodontiformes).* U.S. Department of Commerce, National Oceanic and Atmospheric Administration, National Marine Fisheries Service, Seattle, WA, 1980. doi: 10.5962/bhl.title.63022. URL https://www.biodiversitylibrary.org/item/128327.

[15] J. J. S. Steenstrup and C. F. Lütken. *Spolia atlantica. Bidrag til kundskab om klump- eller maanefiskene (Molidae).* Bianco Lunos Kgl. Hof-Bogrrykkeri (F. Dreyer), Kjøbenhavn, 1898. URL https://www.biodiversitylibrary.org/bibliography/13309.

[25] G. Cuvier and M. Valenciennes. *Histoire naturelle des poissons.* Chez F. G. Levrault, Paris, 1828. URL https://www.biodiversitylibrary.org/page/7111258. Plate 11.

[26] J. Tomelleri. American Fishes, 2019. URL https://www.americanfishes.com/en/content/13-about-contact.

[27] D. S. Jordan, C. W. Metz, and Carnegie Museum. *A catalog of the fishes known from the waters of Korea*. Board of Trustees of the Carnegie Institute, Pittsburgh, 1913. URL https://archive.org/details/catalogoffishesk61jord/page/24/mode/1up.

[28] B. Yau. *Takifugu rubripes*, year of creation unknown. URL https://www.efishalbum.com/details.asp?pid=798.

[29] B. Yau. Fish album terms & conditions, 2022. URL https://www.efishalbum.com/memtermsofuse.asp.

[30] S. J. Raredon. USNM 155442 *Hippocampus erectus* radiograph, year of creation unknown. URL http://n2t.net/ark:/65665/m3cdd9a677-65bc-428e-acee-f55d93aa76bf.

[31] H. M. Smith. *Fishes of North Carolina*. E.M. Uzzell & Co., North Carolina, 1907. URL https://archive.org/details/fishesofnorthcar02smit/page/352/mode/2up.

[32] F. Day. *The fishes of India; being a natural history of the fishes known to inhabit the seas and fresh waters of India, Burma, and Ceylon*, volume 2. B. Quaritch, London, 1878. URL https://www.biodiversitylibrary.org/page/5616157#page/355/mode/1up. Plate CLXIX.

[33] E. Edmonson. Threespine stickleback, 2010. URL https://www.flickr.com/photos/nysdec/29865050421/.

[34] D. Loffler. *Areliscus rhomaleus* RAD107247-001, 1994. URL http://n2t.net/ark:/65665/m3aa836717-9caa-4dba-9318-d64275603924.

[35] B. Yau. *Pseudosciaena crocea*, year of creation unknown. URL https://www.efishalbum.com/details.asp?pid=801.
